# Non-canonical peptidoglycan cross-linking is essential for *Mycobacterium tuberculosis* acid resistance

**DOI:** 10.64898/2026.03.16.712108

**Authors:** Ruoyao Ma, Daniel Rea, Karl Ocius, Ananya Naick, Claire Healy, Thomas R Ioerger, Marcos Pires, Felipe Cava, Sabine Ehrt, Alexandre Gouzy

**Author notes:** Authors contributed equally as co-last authors (Sabine Ehrt, Alexandre Gouzy).

## Abstract

*Mycobacterium tuberculosis* withstands acidic conditions to survive and replicate within macrophages. To define the genetic determinants of this adaptation, we performed a transposon screen in a lipid-rich, acidic medium that mimics the host environment and supports robust *M. tuberculosis* growth. This screen identified *ldtB*, encoding an L,D-transpeptidase, as essential for growth and survival under acid stress. Loss of LdtB decreased 3–3 peptidoglycan cross-linking, disrupted cell wall architecture, and impaired intrabacterial pH homeostasis, resulting in increased susceptibility to cell wall–active antibiotics. Notably, *M. tuberculosis* lacking LdtB displayed heightened sensitivity to meropenem within macrophages, suggesting that targeting this enzyme could potentiate β-lactam efficacy during infection. These findings establish LdtB as a key mediator linking peptidoglycan homeostasis to acid stress resistance and underscores the importance of *in vitro* culture models that recapitulate the host microenvironment for uncovering new *in vivo* active therapeutic targets.

**Teaser:** Disabling a key peptidoglycan cross-linking enzyme compromises *M. tuberculosis* survival under acidic stress and antibiotic exposure.

## Introduction

Tuberculosis (TB), caused by the bacillus *Mycobacterium tuberculosis*, is an ancient disease that has afflicted humans for thousands of years. Evidence of early *M. tuberculosis* infection has been traced to ancient Egyptian mummies and Pre-Columbian Peruvian human skeletons^1,2^. Despite the availability of antibiotics and vaccination, TB remains the leading cause of death from a single infectious agent, responsible for an estimated 10.7 million new cases and 1.23 million deaths worldwide in 2024^3^. As a successful pathogen, *M. tuberculosis* can survive a variety of antimicrobial stresses within its human host^4^. Among these, acid stress is a major host defense strategy and one of the first environmental cues *M. tuberculosis* encounters inside lung macrophages^4–6^. Macrophages contain *M. tuberculosis* in mildly acidic compartments (pH ∼ 6.4) which can reach even lower pH values (pH∼ 4.5) following their activation by lymphocyte-derived interferon gamma (IFN-γ)^7,8^. Resistance to acid stress is therefore a central determinant of *M. tuberculosis* virulence, and mutants defective in acid resistance are attenuated in both macrophage and animal infection models^5,6,8,9^. Understanding how *M. tuberculosis* survives and grows under acidic conditions is thus crucial for elucidating the mechanisms of its pathogenesis and persistence.

Peptidoglycan (PG) is a cell-wall structure essential for bacteria to maintain their shape and protect against environmental threats^10–13^. *M. tuberculosis* PG is a mesh-like heteropolymer consisting of glycan chains of alternating *N*-acetylglucosamine (GlcNAc) and *N*-acetylmuramic acid (MurNAc) or *N*-glycolylmuramic acid (MurNGlyc) residues cross-linked by short peptide stems^14^ (Supplementary Figure 1). Mycobacteria possess two types of PG cross-links, namely 4-3 cross-links and 3-3 cross-links. The classical 4–3 cross-links are generated by D,D-transpeptidases (DDTs), also known as penicillin-binding proteins (PBPs), which use pentapeptide stems as substrates. The non-classical 3–3 PG cross-links are formed by L,D-transpeptidases (LDTs)^15^, which are structurally distinct from DDTs and utilize tetrapeptide stems as substrates^16^ (Supplementary Figure 1). While most bacteria, such as *Escherichia coli* (*E. coli*), rely predominantly on 4–3 cross-linking, mycobacteria are unique in that the majority of their PG cross-links are of the non-canonical 3–3 type throughout all growth phases^17–20^. This observation suggests that LDTs play a major role in *M. tuberculosis* PG homeostasis.

PG maintenance is closely associated with acid stress resistance in bacteria. In *E. coli*, the bifunctional DDT/glycosyltransferase PBP1b is essential for survival under acidic conditions^21^, while the D,D-carboxypeptidase PBP6b is required to maintain rod-shaped morphology in acidic environments^22^. Despite the predominance of 3–3 PG cross-links in *M. tuberculosis*, LDTs have not previously been shown to contribute to acid stress resistance. Notably, prior genetic screens performed with *M. tuberculosis* were conducted under acidic conditions that supported limited growth, potentially hindering the identification of PG pathways essential for cell division^9,23,24^. Our group previously demonstrated that under acidic conditions (pH < 5.8), *M. tuberculosis is* unable to grow on glycolytic carbon sources (e.g., glycerol) but can utilize host-relevant lipids, such as oleic acid (OA) to replicate^23–25^. This improved culture system offers a physiologically relevant platform to dissect mechanisms, such as PG related pathways, that enable *M. tuberculosis* replication in acidic conditions.

Here, we combine functional genomics, biochemical analyses, and macrophage infection models to define the role of the LDT LdtB in *M. tuberculosis* acid resistance. We show that LdtB is essential for maintaining 3–3 cross-linking, PG integrity, and intrabacterial pH homeostasis under acidic conditions. Loss of LdtB sensitizes *M. tuberculosis* to PG-targeting antibiotics, including β-lactams, particularly in acidic conditions *in vitro* and in IFN-γ-activated macrophages. These findings illuminate how *M. tuberculosis* dynamically adapts its cell wall architecture in response to acid stress and reveal stress-specific vulnerabilities that can be exploited for therapeutic intervention.

## Results

### Transposon mutagenesis screening identifies the L,D-transpeptidase LdtB as required for *M. tuberculosis* fitness in acidic conditions

We used transposon mutagenesis coupled with next-generation sequencing (Tn-seq) to identify the genetic determinants important for *M. tuberculosis* replication under acidic conditions. A saturated *M. tuberculosis* transposon library was grown with either glycerol or OA as single carbon source at neutral pH (control conditions) or with OA at acidic pH (test condition) (Fig. 1A). To maximize hit discrimination and enhance statistical sensitivity, the libraries were grown two consecutive times (2 passages) until mid-log phase (OD= 0.5-0.7) before DNA extraction and processing (Supplementary Figure 2). Collected samples showed good sequencing depth and saturation (mean 1.4 million mapped reads per sample and 61.1% saturation), with positive correlations in gene-level insertion counts among biological replicates across all three growth conditions (Pearson r = 0.5–1.0; Supplementary Figure 3).

**Fig. 1.**
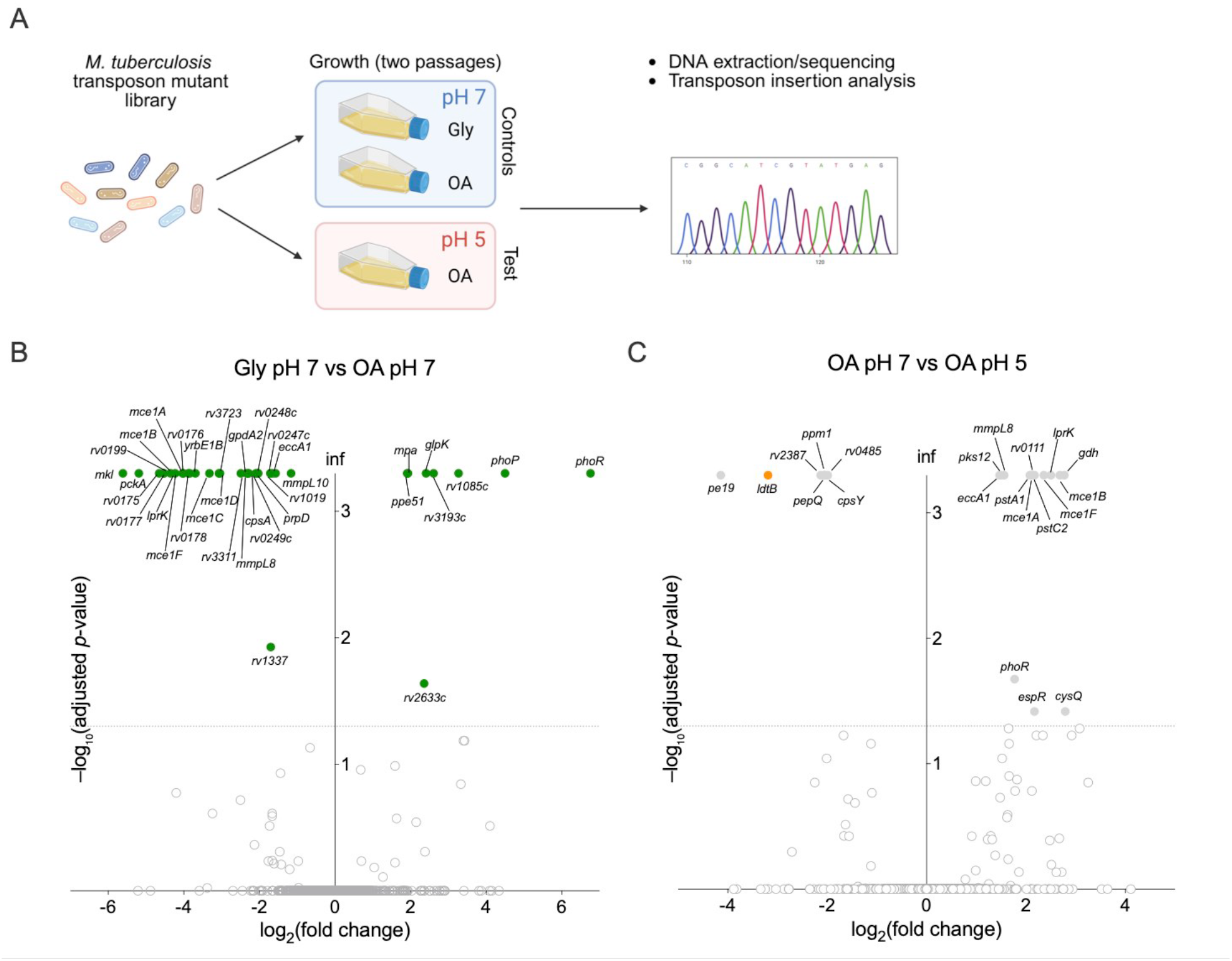
Genome-wide fitness determinants associated with carbon source and acid adaptation. **(A)** Schematic overview of the experimental design for the transposon sequencing (Tn-seq) screen at neutral (pH 7) or acidic pH (pH 5) in the presence of either glycerol (Gly) or oleic acid (OA) as main carbon source. **(B)** Volcano plots showing loci with significantly altered transposon insertion frequencies following two passages in glycerol at pH 7 (Gly pH 7) compared with two passages in oleic acid at pH 7 (OA pH 7). **(C)** Volcano plot showing loci with significantly altered transposon insertion frequencies following two passages in oleic acid at pH 7 (OA pH 7) compared with wo passages in oleic acid at pH 5 (OA pH 5). Genes meeting the statistical threshold (|log₂FC| ≥ 1 and adjusted *p* ≤ 0.05) are shown as colored dots (B, green; C, grey), whereas grey circles represent nonsignificant loci. The *ldtB* gene is highlighted with an orange dot in panel C. Data in panels B and C represent three independent experiments.

Comparative analysis of experiments performed at neutral pH (glycerol pH 7 vs OA pH 7) (Fig. 1B) revealed the importance of genes involved in OA uptake (*mce1* genes, *rv0199*, *lucA*/*rv3723*)^26–29^ and assimilation (*pckA*)^30^ for growth in OA-containing media in comparison to glycerol-containing media (supplemental data 1). As expected, mutants disrupted in genes involved in glycerol uptake (*ppe51*)^31^ and assimilation (*glpK*)^32^ were enriched in OA-containing media, relative to media containing glycerol (Fig. 1B). Notably, mutants interrupted in the two-component system genes *phoP*/*phoR* showed increased fitness in OA-containing media, consistent with the previously reported growth defect of a Δ*phoPR* mutant during growth in glycerol^33^. When comparing the fitness of the libraries depending on pH (Fig. 1C, OA pH7 vs OA pH5), mutants in genes related to phosphate uptake (*pstA1*, *pstC2*) exhibited increased fitness at acidic pH possibly due to their higher capacity to resist acid-mediated oxidative stress^23^. Notably, genes related to OA uptake (e.g. *mce1*, *lprK*) appeared more dispensable in acidic than in neutral conditions (supplemental data 1). This observation could be due to the higher amount of protonated and uncharged OA at acidic pH, which might diffuse through the *M. tuberculosis* plasma membrane independently of the Mce1 uptake system^34^.

Strikingly *ldtB* mutants were among the most under-represented after growth in acidic conditions (Fig. 1C). The *M. tuberculosis* genome encodes five LDTs paralogues that share approximately 30–50% pairwise amino-acid sequence identity with each other^35^, and *in vitro* cross-linking assays demonstrated that four of them, LdtA, LdtB, LdtC and LdtE, possess 3–3 cross-linking activity^36^. Although stress-specific specialization among LDTs has been suggested in *M. tuberculosis*, the physiological roles and environmental contexts in which individual LDTs operate remain largely unexplored. Our data suggest that LdtB might be required for PG 3–3 cross-linking specifically in acidic conditions.

### The L, D transpeptidase LdtB is required by *M. tuberculosis* to grow and survive in acidic conditions

To validate the conditional essentiality of LdtB at acidic pH, we constructed an *ldtB* deletion mutant (Δ*ldtB*) and its corresponding complemented strain expressing *ldtB* under its native promoter (Δ*ldtB*::*ldtB*) (see Materials and Methods and Supplementary Figure 4-6). At neutral pH (pH 7), Δ*ldtB* showed only a minor growth defect relative to WT and the complemented strain, consistent with previous observations (Fig. 2A)^20,37^. Strikingly, at pH 5, Δ*ldtB* ceased replication after ∼10 days and exhibited markedly reduced viability (Fig. 2A, 2B). Complementation restored Δ*ldtB* growth to WT levels, confirming that the observed phenotypes result from *ldtB* loss of function.

**Fig. 2.**
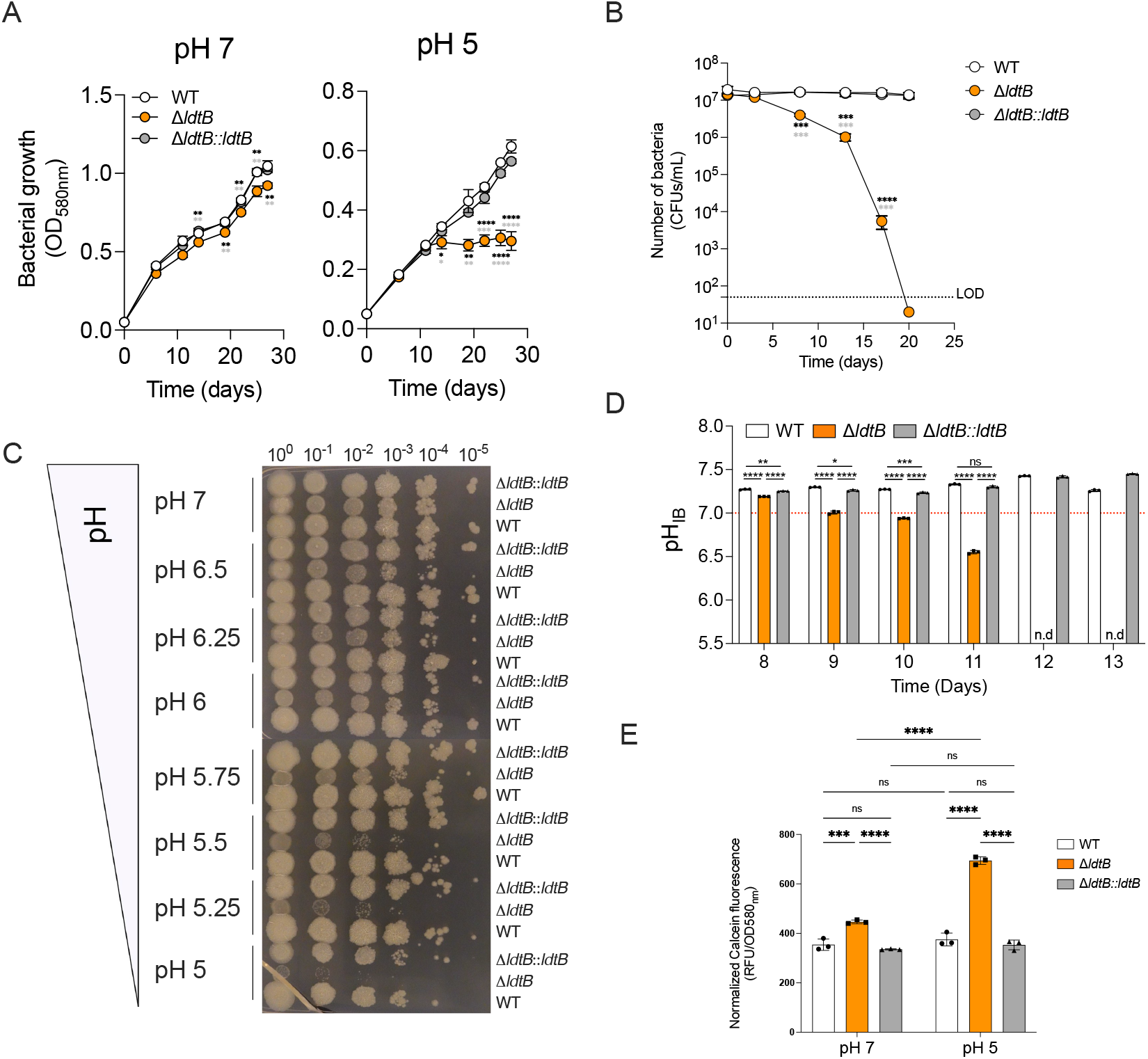
The L,D-transpeptidase LdtB is required for *M. tuberculosis* growth and survival in acidic conditions. **(A)** Growth of WT, Δ*ldtB*, and Δ*ldtB::ldtB* strains in 7H9 medium with oleic acid (OA) at pH 7 or pH 5. Statistical significance was determined using one-way ANOVA with Tukey’s multiple comparisons test; **, adjusted *p* < 0.01; ***, adjusted *p* < 0.001; ****, adjusted *p* < 0.0001. Δ*ldtB* values were compared with either the WT strain (black stars) or the complemented strain (gray stars) **(B)** Survival during prolonged acid exposure. Statistical significance was determined using one-way ANOVA with Tukey’s multiple comparisons test; ***, adjusted *p* < 0.001; ****, adjusted *p* < 0.0001. Δ*ldtB* values were compared with either the WT strain (black stars) or the complemented strain (gray stars). **(C)** Spot assay after 10 days across a pH gradient. **(D)** Measurement of intrabacterial pH (pH_IB_) using the pH-sensitive reporter pHGFP from day 8 to day 13 of incubation in acidic medium (pH 5) supplemented with oleic acid (OA) as main carbon source. The pH_IB_ of the Δ*ldtB* strain carrying pHGFP fell below the limit of detection on days 12 and 13 (n.d = not detectable). Statistical significance was determined using one-way ANOVA with Tukey’s multiple comparisons test; ns, not significant; *, adjusted *p* < 0.05; ***, adjusted *p* < 0.001; ****, adjusted *p* < 0.0001. **(E)** Calcein-AM uptake of WT, Δ*ldtB*, and Δ*ldtB::ldtB* strains cultured at pH 5 and pH 7 for 10 days. Statistical analysis was performed using two-way ANOVA; ns, not significant; ***, adjusted *p* < 0.001; ****, adjusted *p* < 0.0001. Data in panels A to E are representative of three independent experiments.

To further characterize this pH-dependent growth defect, WT, Δ*ldtB* and Δ*ldtB*::*ldtB* strains were cultured across a pH gradient and serial dilutions were spotted onto agar plates at different time points to evaluate bacterial viability. Initial biomasses were comparable among strains at day 0 (Supplementary Figure 7). From day 4 onward, Δ*ldtB* showed decreasing viability starting at pH < 5.75 and worsening at more acidic pH (Fig. 2C & Supplementary Figure 7). These findings demonstrate that LdtB becomes progressively essential during prolonged replication of *M. tuberculosis* under acid stress.

Notably, the expression of both *ldtA* and *ldtB* genes was reported as upregulated in mildly acidic conditions (pH 5.8) suggesting a role for these two LDTs in acid pH adaptation^38^. We thus further examined the impact of pH on the expression of *ldtA*, *ldtB*, along with *ponA1*, a D,D-transpeptidase–encoding gene included as a control. At neutral pH, expression of the PG related genes did not increase over time (Supplementary Figure 8). In contrast, upon exposure to acidic pH (pH 5), *ldtB* expression was rapidly upregulated to a fourfold increase within four hours, whereas *ldtA* expression increased later and was comparatively modest (Supplementary Figure 8). In contrast, *ponA1* expression decreased after four hours in acidic conditions (Supplementary Figure 8). The rapid and specific increase in *ldtB* expression is consistent with the critical role of LdtB for *M. tuberculosis* growth and survival during acid stress that cannot be compensated by LdtA or any other PG-related enzymes.

### LdtB deficiency compromises intrabacterial pH homeostasis and increases cell-envelope permeability in *M. tuberculosis*

*M. tuberculosis* mutants with cell wall defects often exhibit a reduced ability to maintain their intrabacterial pH (pH_IB_) under acid stress^9,39^. To assess whether LdtB contributes to pH_IB_ homeostasis, we monitored pH_IB_ in WT, Δ*ldtB*, and Δ*ldtB*::*ldtB* strains using the ratiometric pH-sensitive GFP (pHGFP) reporter^9,40,41^. When cultured at pH 5, the Δ*ldtB* mutant exhibited a progressive decline in pH_IB_, dropping below 7 by day 10, whereas the WT and complemented strains maintained a near-neutral pH_IB_ (Fig. 2D). These results indicate that LdtB is required for maintaining intracellular pH homeostasis under acidic stress. To determine whether this defect was associated with increased cell envelope permeability, we next performed a calcein-AM uptake assay as previously described^42^. Calcein-AM is a non-fluorescent, membrane-permeable dye that readily enters viable cells and is hydrolyzed by intracellular esterases to calcein, a highly fluorescent molecule that is retained within the cytoplasm and can be readily quantified. On day 10, Δ*ldtB* exhibited significantly greater calcein accumulation than WT and the complemented Δ*ldtB::ldtB* strain at both pH 7 and pH 5, indicating increased cell envelope permeability (Fig. 2E). While calcein accumulation remained pH-independent in WT and Δ*ldtB::ldtB*, Δ*ldtB* showed a significant further increase in permeability at pH 5 compared to pH 7 (Fig. 2E). These results indicate that loss of LdtB function compromises cell envelope integrity and that this defect is exacerbated during prolonged exposure to acidic conditions.

The increased cell-envelope permeability of Δ*ldtB* during prolonged acid exposure coincided with greater intrabacterial acidification (Fig. 2D) and reduced viability (Fig. 2B). Together, these findings support a role for LdtB-mediated 3–3 PG cross-linking in maintaining cell envelope integrity, intrabacterial pH homeostasis, and bacterial survival under acidic conditions.

### Loss of LdtB impairs 3–3 cross-linking, shortens cells, and disrupts cell-wall integrity under acidic conditions

To examine pH-dependent 3-3 cross-linking by LdtB, we measured the incorporation of TetraFl, a fluorescent tetrapeptide probe that reports overall LDT activity^43^. *M. tuberculosis* strains were incubated overnight with TetraFl in neutral (pH 7) or acidic media (pH 5), followed by fluorescence microscopy and quantification by flow cytometry. Microscopy revealed reduced labeling of Δ*ldtB* cells at pH 7 and markedly more so at pH 5 relative to the WT strain (Fig. 3A). Quantitative analysis by flow cytometry showed that the mean fluorescence intensity (MFI) of Δ*ldtB* cells was approximately 2.32-fold lower than that of WT at pH 7 and 5.6-fold lower at pH 5 (Fig. 3B, Supplementary Figure 9 and Supplementary Figure 10). These findings demonstrate that LdtB performs 3–3 crosslinking *in vivo* with minor activity at neutral pH and is the predominant LDT under acidic conditions.

**Fig. 3.**
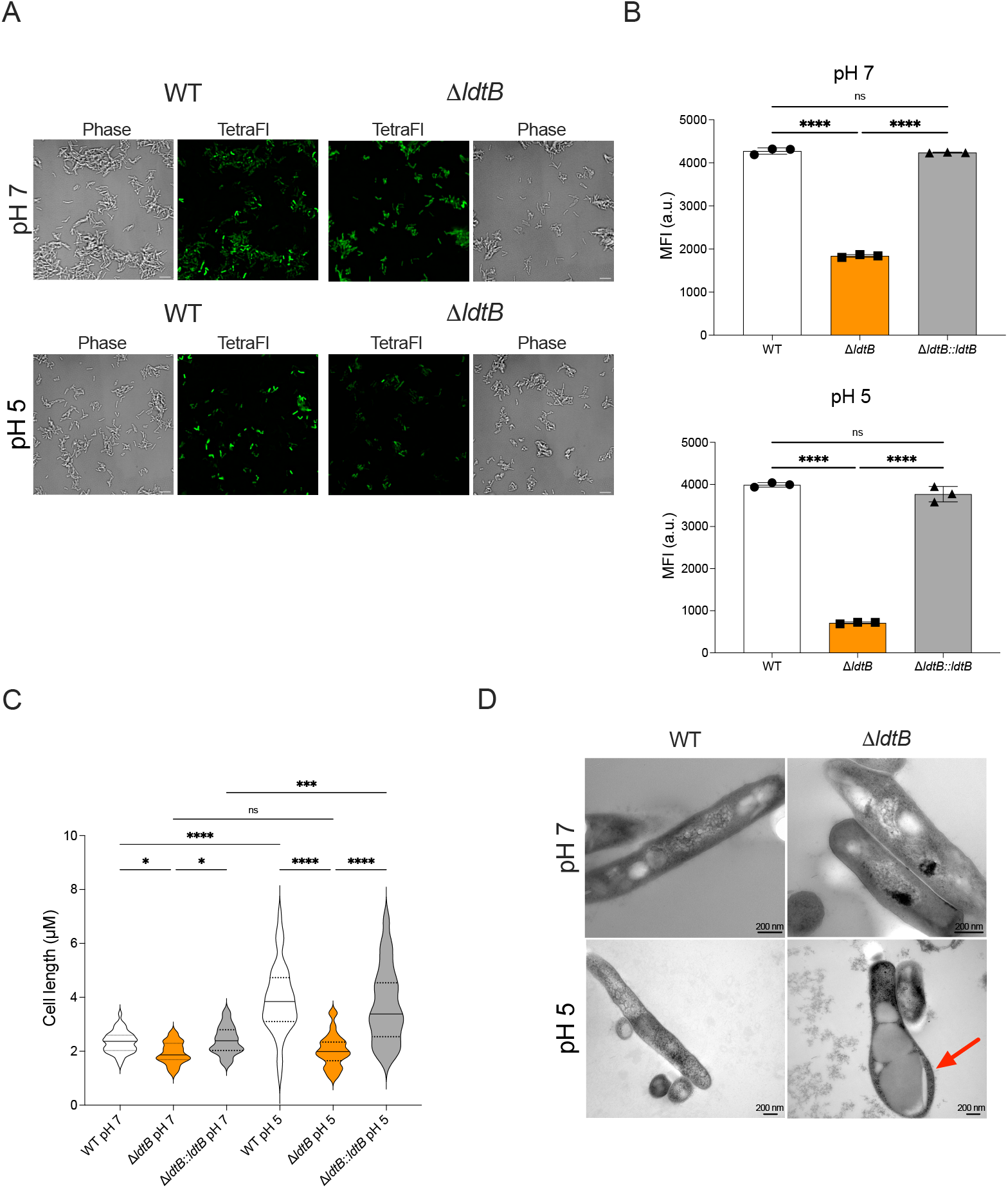
Loss of LdtB decreases tetrapeptide incorporation into peptidoglycan (PG) and disrupts *M. tuberculosis* cell wall under acidic conditions. **(A)** Fluorescence and phase-contrast micrographs of WT and Δ*ldtB* cells labeled with TetraFl overnight at pH 7 or pH 5 in medium containing oleic acid (OA) as main carbon source. Scale bar, 5 μm. **(B)** Flow cytometry analysis of TetraFl incorporation on the indicated strains. Data are representative of three independent experiments. **(C)** Quantification of cell length for WT, Δ*ldtB*, and Δ*ldtB::ldtB* strains (n = 47) after 8 days at pH 5 or pH 7; statistical significance was determined using the Kruskal–Wallis test with post-hoc multiple comparisons; ns, not significant; *, adjusted *p* < 0.05; ****, adjusted *p* < 0.0001. Data are representative of three independent experiments. **(D)** Representative transmission electron micrographs of WT and Δ*ldtB* cells cultured for 8 days at pH 7 or pH 5. The red arrow indicates a bulging region in the Δ*ldtB* mutant at pH 5. Images were acquired at an accelerating voltage of 100 kV and nominal magnification was ×80,000 (WT, pH7), ×100,000 (Δ*ldtB*, pH7) and ×50,000 (WT, pH5; Δ*ldtB,* pH5). Scale bars, 200 nm.

Prior work demonstrated that Δ*ldtB* cells are shorter than WT at neutral pH^37,44^. Our microscopy data support this observation and further show that Δ*ldtB* cells are also substantially shorter at acidic pH (Fig. 3C). Remarkably, WT *M. tuberculosis* bacilli increased in length during prolonged acid exposure (day 8 Fig. 3C). While the cellular mechanisms governing this acid-induced cell elongation remain to be elucidated, we show that this process requires LdtB-mediated 3–3 peptidoglycan cross-linking.

Quantification of cells with bulged or irregular morphology revealed that approximately 24% of Δ*ldtB* cells exhibited abnormal morphology at pH 5 following prolonged acid exposure (day 8; Supplementary Figure 11). To further evaluate structural consequences of LdtB deficiency, we then performed transmission electron microscopy (TEM) on Δ*ldtB* at neutral and acidic pH. Pronounced cell envelope abnormalities were observed specifically in Δ*ldtB* cells under acidic conditions, including cellular bulging, ghost cells lacking intact cell wall layers, and localized cell envelope defects (Fig. 3D & Supplementary Figure 12). These findings indicate that under acid stress *M. tuberculosis* requires LdtB to maintain PG integrity and prevent structural breaches that ultimately cause cell lysis.

### PG analysis reveals 3-3 crosslinking activity is predominantly mediated by LdtB in acidic conditions

To biochemically define LdtB’s role in PG homeostasis, we analyzed PG composition from WT, Δ*ldtB* and Δ*ldtB*::*ldtB* strains depending on pH using ultra-performance liquid chromatography-mass spectrometry (UPLC-MS) (Fig. 4). Schematic representations and nomenclature of monomeric and dimeric PG structures are shown in Supplementary Figure 6. Analysis of PG fragment abundance in the WT strain at pH7 (Fig 4 A & C) and pH 5 (Fig 4 B & D) revealed that the ratio of 4–3/3–3 cross-links remained stable across pH conditions (Fig. 4E and F) consistent with previous observations in non-replicating conditions^19^. This suggests that the 4–3/3–3 cross-link ratio is a defining feature of the *M. tuberculosis* cell wall. At pH 7, Δ*ldtB* exhibited a minor but statistically significant reduction (∼14%) in 3–3 cross-links relative to WT and complemented strains, with no significant changes in 4–3 cross-links (Fig. 4E). Strikingly, Δ*ldtB* showed a much more pronounced reduction (∼60%) in 3–3 cross-links pH than WT and complemented strains at acidic (Fig. 4F). Accordingly, the tetrapeptide monomer (M4_Mt_ peak; Fig. 4A-D; Supplementary Figure 13) which serves as donor muropeptide substrate for 3-3 crosslinks was more abundant in Δ*ldtB* than in WT and complemented strains (Fig 4. A-D). Notably, 4–3 cross-linking in Δ*ldtB* was slightly elevated at acidic pH (statistically significant only relative to the complemented strain) (Fig. 4F), suggesting a compensatory DDT activity to preserve PG cross-linking in the absence of LdtB.

**Fig. 4.**
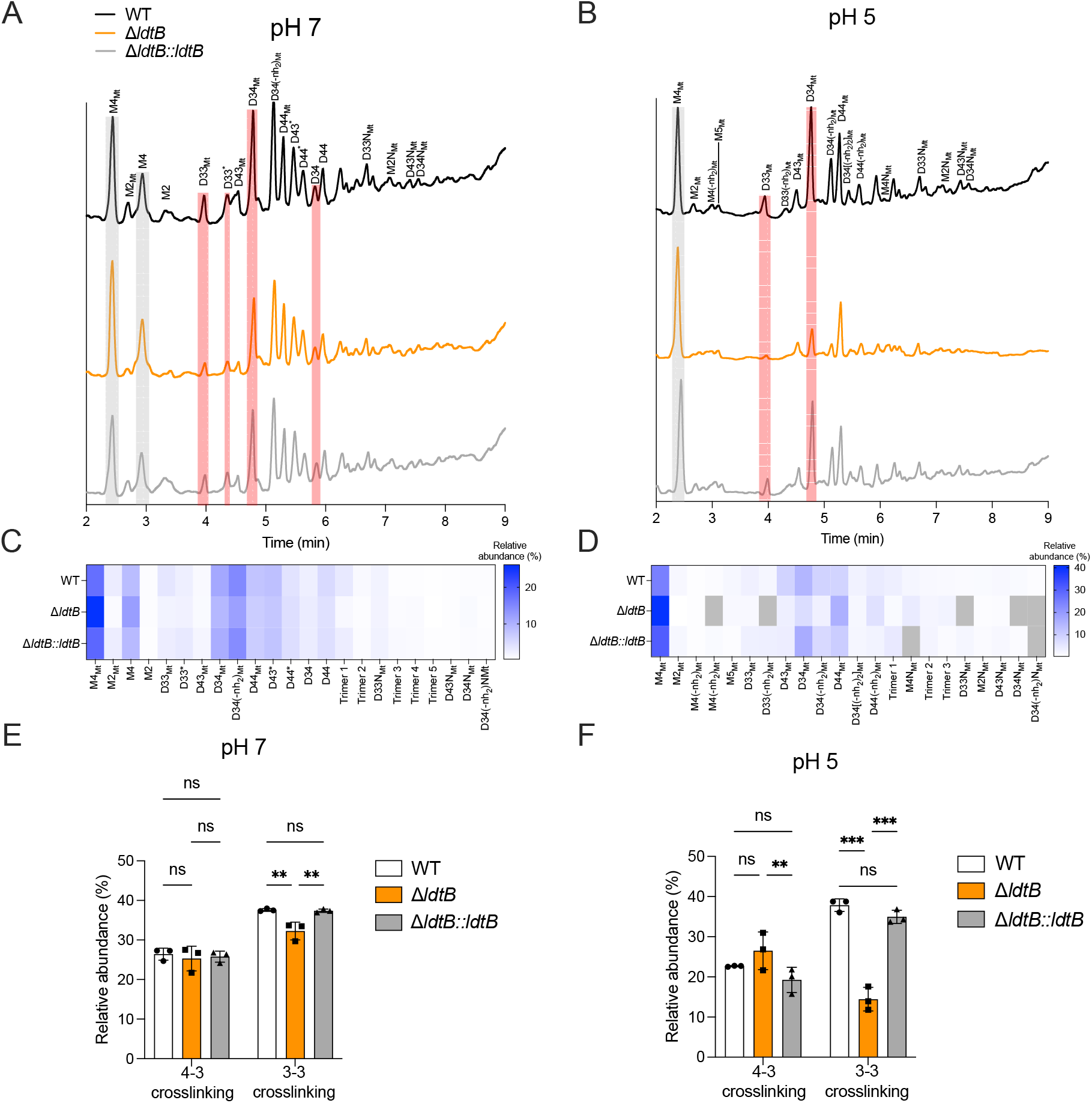
LdtB promotes 3–3 peptidoglycan cross-link formation in *M. tuberculosis* under acidic stress. (A-B) Representative mass chromatograms showing ionization profiles of peptidoglycan from WT, Δ*ldtB*, and Δ*ldtB::ldtB* strains grown under neutral **(A)** or acidic **(B)** conditions. Chromatograms shown are representative of three biological replicates. **(C-D)** Heatmaps depicting the relative abundance of muropeptides detected in A-B under neutral **(C)** or acidic **(D)** conditions. The color scale indicates relative abundance from white (low) to blue (high); grey indicates muropeptides that were not detected. **(E-F)** Quantification of 4–3 and 3–3 cross-links in WT, Δ*ldtB*, and Δ*ldtB::ldtB* under neutral (E) and acidic (F) conditions. Statistical analysis was performed using two-way ANOVA; ns, not significant; **, adjusted *p* < 0.01; ***, adjusted *p* < 0.001. Data in panels E and F are from three biological replicates and are presented as mean ± SD.

Remarkably, PG isolated from strains grown at pH 5 displayed increased levels of *N*-glycolyl muramic acid residues. Fully glycolylated monomers and dimers were detected at pH 5 (“Mt” peaks), whereas PG from strains grown at pH 7 showed mixed acetylated and glycolylated dimers (D33*, D43*, and D44* peaks) or exclusively acetylated species (M4, M2, D34 and D44) (Fig. 4A-D & Supplementary Figure 13). Previous studies suggested that *N*-glycolylation provides stronger structural stability to the cell wall by enhancing hydrogen bonding resulting in tighter binding of the PG^14,46^. Altogether, these results demonstrate that LdtB is essential for maintaining 3–3 cross-links under host-relevant acidic conditions and that *M. tuberculosis* increases PG glycolylation at acidic pH possibly to strengthen PG structure.

### LdtB protects *M. tuberculosis* from PG-targeting antibiotics under acidic conditions

Because LdtB is required for proper PG cross-linking and maintenance of cell wall integrity under acidic conditions, we hypothesized that *ldtB* deletion would increase susceptibility to PG-targeting antibiotics. We measured the minimum inhibitory concentrations (MICs) of representative PG-targeting antibiotics in WT, Δ*ldtB*, and Δ*ldtB*::*ldtB* grown at pH 7 or pH 5.75. We used pH 5.75 (instead of pH 5.0) to permit sufficient growth of Δ*ldtB* for growth inhibition assays (Fig. 2C). The PG synthesis inhibitors tested included β-lactams, meropenem and amoxicillin clavulanate, as well as the glycopeptide vancomycin (Supplementary Figure 1), with the non-PG targeting drug isoniazid (INH) as a control.

At neutral pH (pH 7), Δ*ldtB* showed slight (amoxicillin clavulanate) to moderate (meropenem; vancomycin) increases in susceptibility to the PG targeting drugs compared to WT and complemented strains (Fig. 5A) consistent with previous studies^20,37,44^. This observation aligns with LdtB’s minor role in PG 3–3 cross-linking and *M. tuberculosis* growth at pH 7 (Fig. 2A, 3A-3B & 4). In contrast, at acidic pH, Δ*ldtB* displayed a markedly increased sensitivity to meropenem, vancomycin, and amoxicillin-clavulanate (Fig. 5B). These findings demonstrate that the reduced 3-3 crosslinking in Δ*ldtB* at acidic pH renders the mutants hypersusceptible to PG-targeting agents. Importantly, susceptibility to the non-PG-targeting antibiotic INH was unchanged in the absence of LdtB, indicating that increased Δ*ldtB* susceptibility to PG-targeting agents is not due to generally increased drug permeability. Together, these results demonstrate that LdtB activity protects *M. tuberculosis* against PG-targeting antibiotics, especially under acidic conditions.

**Fig. 5.**
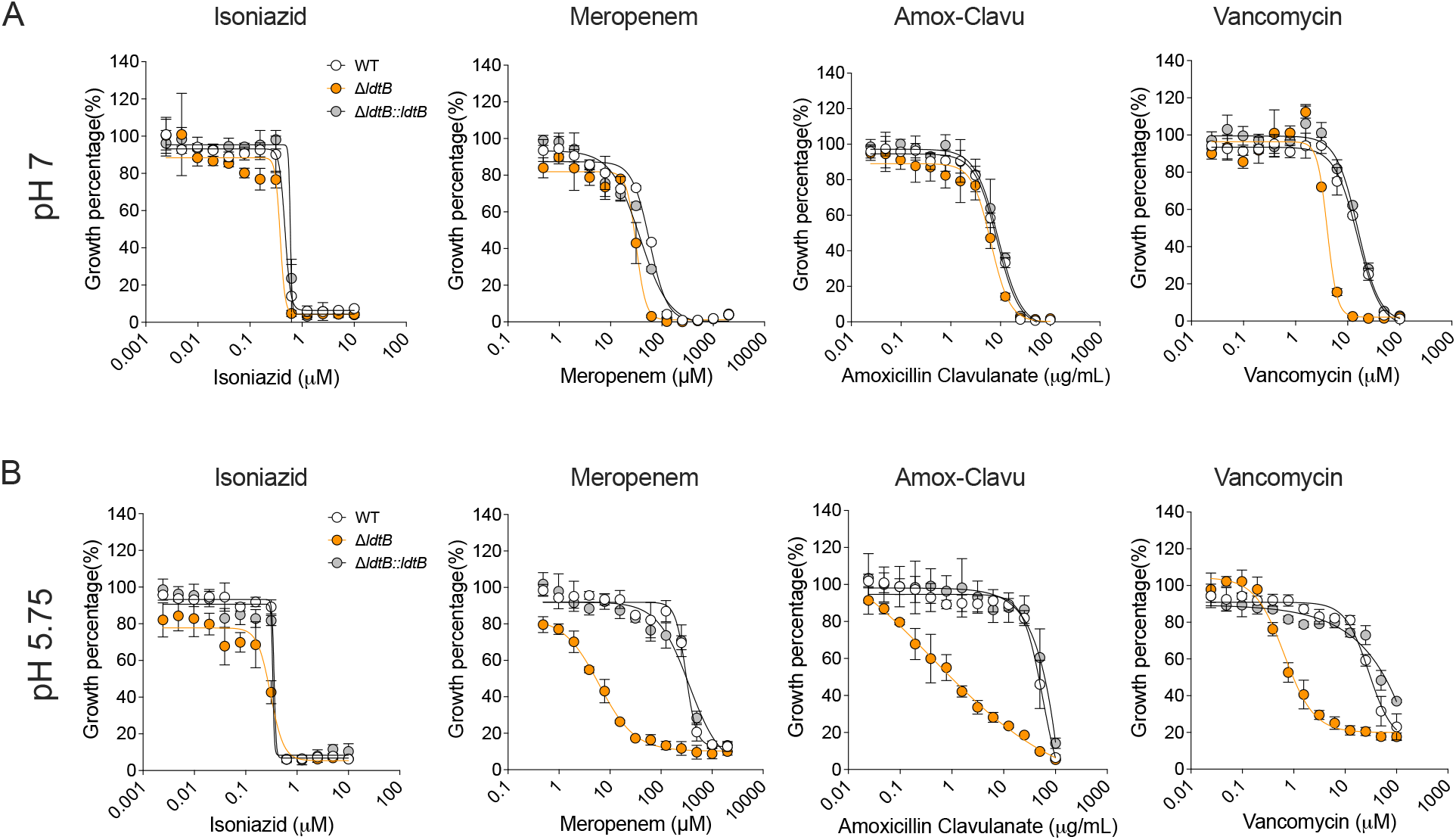
Loss of LdtB sensitizes *M. tuberculosis* to peptidoglycan-targeting antibiotics in acidic conditions *in vitro*. Comparison of MIC values for isoniazid, meropenem, amoxicillin clavulanate, and vancomycin against WT, Δ*ldtB*, and Δ*ldtB*::*ldtB* strains in neutral (pH 7) **(A)** and acidic (pH 5.75) **(B)** media. Data are means of three technical replicates and are representative of three independent experiments.

### LdtB protects *M. tuberculosis* against meropenem during macrophage infection

Next, we investigated the role of LdtB in protecting against PG-targeting antibiotics in a more physiological model, the macrophage. We first tested the fitness of Δ*ldtB* in resting and IFN-γ-activated murine bone marrow-derived macrophages (BMDMs) in the absence of any drugs. We observed no significant replication defect of Δ*ldtB* compared to WT (Supplemental Figure 14) consistent with the Δ*ldtB* growth emerging only during prolonged replication at acidic pH (≥ day 10, Fig. 2A), a condition not recapitulated in the *ex vivo* macrophage model. To better reveal Δ*ldtB* susceptibility to PG-targeting drugs, we therefore preconditioned *M. tuberculosis* strains for 3 days in medium at pH 7 or pH 5 before infecting resting and IFN-γ-activated BMDMs, then treated macrophages with meropenem (100 or 1000 µM) or left untreated for 5 days before quantifying bacterial survival.

For bacteria preconditioned at pH 7, meropenem treatment caused a dose-dependent reduction in bacterial viability that was similar for WT, Δ*ldtB*, and complemented strains in both resting and IFN-γ-activated macrophages (Fig. 6A & 6B, left panels & Supplementary Figures 8A & B, left panels). In contrast, after preconditioning at pH 5, Δ*ldtB* exhibited a moderate but significant reduction in CFUs upon meropenem treatment in resting macrophages compared to WT, though not statistically significant relative to the complemented strain (Supplementary Figures 15A & B, right panels). Strikingly, in IFN-γ-activated macrophages infected with acid-preconditioned bacteria, Δ*ldtB* showed a concentration dependent and significant decrease in viability compared with WT and complemented strains upon meropenem treatment (Fig. 6A & 6B, right panels). These results demonstrate that LdtB protects *M. tuberculosis* from meropenem during macrophage infection particularly acid exposed bacteria within IFN-γ-activated macrophages. The data are consistent with the activity of IFN-γ in acidifying *M. tuberculosis*-containing intracellular compartments, which we confirmed using two complementary microscopy approaches (Supplementary Figure 16A-C)^7–9^, and reveal that loss of LdtB-dependent PG 3–3 crosslinking can potentiate β-lactam activity against *M. tuberculosis* during macrophage infection.

**Fig. 6.**
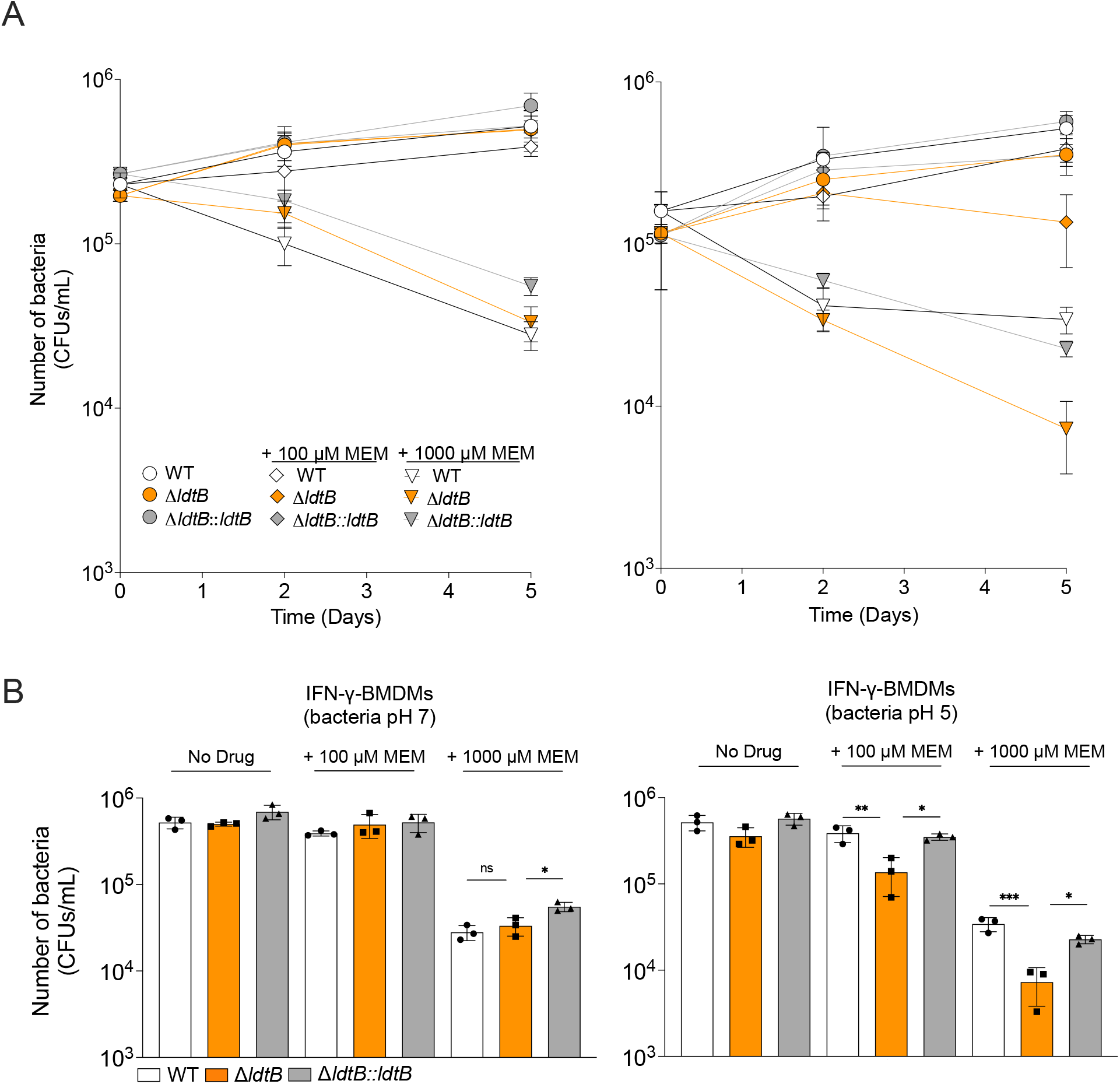
LdtB protects *M. tuberculosis* against meropenem within IFN-γ activated macrophages. **(A)** CFU of WT, Δ*ldtB*, and Δ*ldtB::ldtB* strains in IFN-γ activated bone marrow–derived macrophages preconditioned in neutral (left panel) or acidic media (right panel) and treated with increasing concentrations of meropenem. **(B)** Quantification of CFU on day 5 from panel A. Statistical significance was determined using one-way ANOVA with Tukey’s multiple comparisons test; ns, not significant; **, adjusted *p* < 0.01; ***, adjusted *p* < 0.001. CFU data represent mean ± SD from three technical replicates per condition and are representative of three independent experiments.

## Discussion

*M. tuberculosis* is a successful pathogen, in part because it survives within hostile environments such as the acidic compartments within macrophages^4,47–49^. Our study provides mechanistic insight into how the L, D-transpeptidase LdtB enables *M. tuberculosis* to maintain cell wall integrity and resist acid stress. We demonstrate that in acidic conditions, *M. tuberculosis* relies predominantly on LdtB-mediated 3–3 PG cross-linking to preserve cell wall integrity. Despite the inherent difficulty of PG composition analysis in *M. tuberculosis*, we applied UPLC-MS to define PG chemical composition after acid exposure and found that high 3–3 cross-linking (∼40% relative abundance) is required for *M. tuberculosis* to withstand acid stress, whereas reduced 3-3 cross-linkage (∼20% relative abundance in Δ*ldtB*) leads to impaired growth, reduced intrabacterial pH and compromised PG integrity. Consequently, loss of LdtB markedly increases susceptibility to β-lactam antibiotics, such as meropenem, particularly in acidic conditions and during infection of IFN-γ-activated macrophages.

Our findings are consistent with the reported role of LdtB in *M. tuberculosis* virulence in mice^20^. An Δ*ldtB* mutant established infection and grew similarly to WT during the first two weeks of infection but failed to persist after the onset of adaptive immunity. This pattern was observed for other *M. tuberculosis* acid-susceptible mutants^6,9^ and likely reflects increased acid stress within infected IFN-γ-activated macrophages during chronic infection^7^.

Although *M. tuberculosis* encodes multiple LDTs, our results indicate that they are not functionally redundant under specific stresses. In standard growth conditions, LdtC, may compensate for the loss of LdtB activity as previously suggested^37^. In accordance, structural comparisons revealed that LdtB and LdtC are the most similar LDTs in *M. tuberculosis*^50^. However, under acid stress, reduced activity or expression of LdtC and the other LDTs appears to prevent effective compensation, rendering LdtB essential for viability. Future studies measuring the expression, stability and enzymatic activity of LDTs under varying environmental conditions will help identify the dedicated roles of individual enzymes.

Altogether, our data support a model in which *M. tuberculosis* survival in acidic environments relies on the rewiring of its PG machinery. In line with this model, the serine protease MarP regulates the activity of the PG endopeptidase RipA and is required for pH_IB_ homeostasis and survival in acidic conditions^39,51,52^. Future studies should determine how MarP/RipA and LdtB control PG remodeling and cell division in acid-stressed *M. tuberculosis*. Notably, *Salmonella enterica* serovar Typhimurium, also preferentially utilizes specific virulence-associated DDTs to grow under acidic conditions^53,54^. Our study reinforces the concept that PG-mediated acid stress resistance is a common trait necessary for virulence of intracellular pathogens such as *M. tuberculosis* and *S. enterica*.

Using our lipid rich and acidic growth medium, we further show that *M. tuberculosis* growth in acidic conditions is associated with a substantial increase in cell length (Fig. 3C). Previous studies did not observe this, possibly because the culture media used failed to support robust replication, thereby masking this pH-dependent morphological trait. Increased cell length has also been observed in clinical *M. tuberculosis* isolates replicating inside macrophages^55^. Our data suggest that acidic pH is responsible for the cell elongation during infection. The trigger(s) and functional consequences of this pH-associated cell elongation in *M. tuberculosis* pathogenesis remain to be defined.

In addition to altering cell length, our findings indicate that environmental pH shapes PG modification in *M. tuberculosis*, particularly *N*-glycolylation, a hallmark of Actinomycetes cell wall structure. Studies in standard media reported that *M. tuberculosis* PG contains a mixture of *N*-acetylated and *N*-glycolylated muramic acid residues^46,56–58^ and we similarly detected both species at neutral pH; however, in acidic conditions, *M. tuberculosis* produced exclusively *N*-glycolylated muropeptides perhaps the result of enhanced activity of the *N*-acetyl muramic acid hydroxylase Rv3818/NamH, mediating PG glycolylation^14^. Increased PG glycolylation likely protects *M. tuberculosis* from lysozyme-mediated lysis in acidic intracellular compartments^14,59^ and may also augment host recognition via NOD2 signaling, ultimately leading to increased TNF production by macrophages^14,60–62^. Similarly, other intracellular pathogens such as *S. enterica* edit their PG to modulate their recognition by host cells^63^. Taken together, our data indicate that pH-dependent PG alterations, such as *N*-glycolylation, allow *M. tuberculosis* to maintain cell wall integrity and modulate host-pathogen interactions.

From a therapeutic standpoint, our work reveals an exploitable vulnerability in *M. tuberculosis*’s acid stress response. Although β-lactam antibiotics are widely used, they have limited activity against *M. tuberculosis* in part because of the beta-lactamase BlaC^64,65^. Carbapenems such as meropenem, which target both DDTs and LDTs^45^, resist BlaC-mediated inactivation and have gained attention for their efficacy in treating multidrug-resistant and extensively drug-resistant TB (MDR/XDR-TB)^66,67^. Together with the recent identification of specific LdtB inhibitors^69^, these observations provide a rationale for future studies to determine whether pharmacological inhibition of LdtB can enhance β-lactam activity. Further synergy studies, pharmacodynamic analyses, and *in vivo* treatment experiments will be required to establish the therapeutic relevance of targeting LdtB. More broadly, our study establishes a link between acid resistance, PG cross-linking, and antibiotic susceptibility in *M. tuberculosis* and highlights stress-dependent cell wall remodeling as a potential area for further investigation.

## Materials and Methods

### Culture conditions

Mycobacterial cultures were cultured in Middlebrook 7H9 broth supplemented with 0.2% glycerol, 0.05% tyloxapol, and ADN (0.5% bovine serum albumin [BSA], 0.2% dextrose, and 0.085% NaCl), or on Middlebrook 7H10 agar containing 0.5% glycerol and 10% oleic acid-albumin-dextrose-catalase (OADC) enrichment (Becton Dickinson). For pH-dependent growth assays, fatty acid-free (FAF) 7H9 medium was prepared as previously described and supplemented with 200 μM oleic acid (OA) every 2-3 days^25^. For MIC assays and calcein-AM uptake asssays, media formulations containing high BSA and OA, FAF-BSA was used at 50 g/L and OA was added to a final concentration of 3 mM. For intrabacterial pH measurement assays, Sauton’s “No Carbon” medium was prepared containing 0.05% (w/v) KH₂PO₄, 0.2% (w/v) citric acid, 0.05% (w/v) (NH₄)₂SO₄, 0.005% (w/v) ferric ammonium citrate, 0.001% (v/v) ZnSO₄ (from a 1% stock solution) and 0.05% (v/v) tyloxapol and 10% (v/v) AN solution (0.5% FAF-BSA and 0.085% NaCl). This Sautons media was then supplemented with 200 μM OA every 2-3 days during experiments. The pH of the media was adjusted using HCl or NaOH as appropriate. MES (2-(N-morpholino)ethanesulfonic acid) or MOPS (3-(N-morpholino)propanesulfonic acid) was added to a final concentration of 100 mM to stabilize acidic or neutral pH, respectively. When required, antibiotics were used at the following final concentrations: hygromycin, 50 μg/mL; kanamycin, 25 μg/mL; streptomycin, 25 μg/mL.

### Construction of the Mycobacterium tuberculosis ΔldtB mutant

The *M. tuberculosis* Δ*ldtB* mutant was generated in the H37Rv background using a RecET-mediated recombineering approach. Briefly, *M. tuberculosis* H37Rv carrying the pNit-RecET-sacB-kanR plasmid was grown to mid-log phase, and expression of the RecET proteins was induced by addition of 10 μM isovaleronitrile. After 8 h of induction, the culture was treated with 2M glycine and incubated for an additional 16 h. A DNA fragment containing a hygromycin resistance cassette flanked by 500 bp homologous to the regions upstream and downstream of *ldtB* was synthesized (GenScript Biotech) and introduced by electroporation. Recombinants were selected on 7H10 agar supplemented with hygromycin. Spontaneous loss of the pNit-RecET-sacB-kanR plasmid and deletion of *ldtB* were confirmed by PCR (Supplementary Figure 4). The resulting Δ*ldtB* mutant was verified by whole-genome sequencing (SeqCenter) to confirm deletion of *ldtB*, assess secondary mutations, and identify phthiocerol dimycocerosate (PDIM)-positive clones (Supplementary Figure 5). Sequencing reads were aligned to the *M. tuberculosis* H37Rv reference genome using BWA-MEM^68^, and aligned files were processed using SAMtools (Picard Toolkit) and Picard^69^. Variants were called using GATK^70^, and PDIM status was assessed using an in-house Python script incorporating Biopython^71^. Sequencing coverage across the *ldtB* locus was visualized using pyGenomeTracks^72^ to confirm the deletion. For genetic complementation, the verified Δ*ldtB* mutant was transformed with an integrative plasmid expressing *ldtB* under the control of its native promoter (a 138 bp region upstream of the start codon) integrated at the attL5 site. The expression of *ldtB* in WT, Δ*ldtB* and Δ*ldtB::ldtB* was assessed by RT-qPCR (Supplementary Figure 6).

### Transposon mutagenesis coupled with next-generation sequencing (Tn-seq)

An *M. tuberculosis* H37Rv saturated transposon library (himar1 transposon) previously constructed was used^23^. A frozen aliquot of the transposon mutant library was cultured for 3 days in 25 mL standard 7H9 media containing 10X ADN, tyloxapol 0.05%, glycerol 0.2% (∼ 22mM) and kanamycin 25 µg/mL. Bacteria were then washed two times and resuspended using PBS-tyloxapol 0.05%. Bacteria were inoculated in media at a starting OD of 0.05. Three types of media were used. Neutral pH media containing glycerol (glycerol 0.2%) as main carbon source, neutral pH media containing oleic acid (OA) (200µM) as main carbon source and acidic pH media (pH 5.0) containing oleic acid (OA) (200µM) as main carbon source. 100mL of each media was used for each condition and pH was adjusted by addition of HCl/NaOH using a pH meter before inoculation. When OA was the main carbon source used, 200µM of OA was supplemented to the cultures every 2-3 days. Fatty acid free BSA (0.5%) and 0.085% NaCl (FAF-AN) was used to supplement the defined carbon media. Bacteria from the first passage were collected when the OD of the cultures reached between 0.5-0.7 and used to inoculate fresh media for a second passage. The bacteria from the second passage were also collected at OD between 0.5-0.7. At the end of each passage, genomic DNA was extracted from the bacterial pellets and the library mutant composition was determined by sequencing amplicons of the transposon-genome junctions as described previously^23^. Mapping and quantification of transposon insertions was done as described previously using the TRANSIT Tn-seq analysis tool^23,73,74^ to identify transposon mutants that were under-or over-represented (using the resampling algorithm for pairwise comparisons; log2 fold change, greater than 1 or less than −1) with statistical significance after correction for multiple comparisons (adjusted *p* value ≤ 0.05). The screen was repeated three times by performing three independent experiments. Triplicates were analyzed jointly using TRANSIT. Briefly, reads were mapped to the H37Rv genome, generating wig files (with insertion counts at TA sites) for the 9 samples (three replicates for the three conditions tested). The statistics of the samples are shown in Supplementary Table 1. The number of mapped reads was 1.4 million read per sample on average (range 0.61-2.0M). The saturation of each sample was good, with a mean of 61.1% (range: 57-66%). The mean insertion count at non-zero TA sites was 22.4. Supplementary Figure 3A shows a correlation plot of the 9 samples. The correlation coefficient (Pearson *r*) of gene means between most pairs of samples was good (between 0.5 and 1.0). Supplementary Figure 3B, 3C and 3D show the pairwise scatterplots of gene means between pairs of replicate samples for growth on glycerol at pH 7, oleic acid pH7 and oleic acid pH5, respectively.

### Transmission electron microscopy (TEM) and wide-field light microscopy

*M. tuberculosis* WT and Δ*ldtB* strains were grown to mid-log phase in standard Middlebrook 7H9 medium, harvested by centrifugation, and washed twice with PBS-tyloxapol 0.05%. Bacterial were diluted to an OD_580nm_ of 0.05 in 7H9 HBO medium adjusted to either pH 5 or pH 7 and cultured for 8 days, reaching OD_580nm_ values of 0.3-0.4 for the pH 5 cultures and 0.8-1.0 for the pH 7 cultures. Mycobacterial pellets were fixed in 1.25% glutaraldehyde, 4% paraformaldehyde, 4% sucrose, 0.075% L-lysine, 0.01 M CaCl₂ and 0.01 M MgCl₂ in 0.1 M sodium cacodylate buffer (pH 7.3) for ≥4 h and then overnight at 4°C. Samples were washed three times in cacodylate buffer and post-fixed for 1 h in 1% osmium tetroxide, 4% sucrose and 1% tannic acid. Pellets were washed, dehydrated through a graded ethanol series, and infiltrated with Quetol/Spurr’s resin, first at a 1:1 ethanol:resin ratio for 30 min and then in pure resin overnight. Samples were embedded in fresh resin and polymerized at 50°C for 24 h. Samples were then sectioned to 65–70 nm using a diamond knife (Diatome) on a Leica Ultracut UCT ultramicrotome. All imaging was performed using a transmission electron microscope (JEM-1400; JEOL Ltd.) at Weill Cornell Medical College. For cell length measurements and quantification of abnormal cells, cultures harvested under the conditions described above were washed three times with PBS-tyloxapol 0.05% and fixed with 4% paraformaldehyde (PFA) for 4 h at room temperature before removal from BSL-3 containment. PFA-Fixed bacterial suspensions were immobilized on 7H9 pads containing 1% (w/v) agarose and imaged using an inverted IX-70 microscope (Olympus) equipped with an sCMOS camera (pco.edge). Image analyses were performed using ImageJ software^75^.

### RNA extraction and RT-qPCR

20 mL of Mycobacterial culture were mixed with an equal volume of guanidinium thiocyanate buffer and centrifuged for 10 minutes at 4 °C. The bacterial pellets were resuspended in 1 mL TRIzol reagent and lysed by bead beating. Beads were removed by centrifugation, and the resulting supernatants were transferred to new tubes. 200 μL of chloroform were added, followed by vigorous mixing. RNA was purified using the RNA Clean & Concentrator-5 Kit (Zymo Research), and gDNA was removed with TURBO DNase (Ambion). Samples were subsequently repurified using the RNA Clean & Concentrator-25 Kit (Zymo Research). Complementary DNA (cDNA) was synthesized using MuLV Reverse Transcriptase (New England Biolabs) in a 25 µL reaction. A reaction lacking both the RNase inhibitor and MuLV enzyme was included to confirm the absence of genomic DNA contamination. Quantitative PCR was performed using the LightCycler® 480 System (Roche). The housekeeping gene *sigA* (*Rv2703*) served as an internal control. Primers and probes used for RT-qPCR are listed in the Supplementary Table 2.

### PG labeling probe synthesis and fluorescence microscopy

PG labeling probes (TetraFl) were synthesized as previously described^43,76^. For PG labeling, 100 µM TetraFl was added to *M. tuberculosis* cultures at the indicated time points. Samples were collected, washed three times with PBS-tyloxapol 0.05%, and fixed with 4% paraformaldehyde (PFA) for 4 h at room temperature before removal from BSL-3 containment. PFA-Fixed bacterial suspensions were immobilized on 7H9 pads containing 1% (w/v) agarose and imaged using an inverted IX-70 microscope (Olympus) equipped with an appropriate fluorescence filter set, a sCMOS camera (pco.edge), and an Insight SSI 7-color solid-state illumination system. Image analyses were performed using ImageJ software^75^.

### Flow cytometry analysis of mycobacteria labeling with TetraFl

PFA-fixed *M. tuberculosis* samples were washed twice with PBS-tyloxapol 0.05% and resuspended in PBS. Flow cytometric analysis was performed on a LSRFortessa X-20 cell analyzer (BD Biosciences) using the 488 nm blue laser and the 530/30 nm bandpass (FITC) detector for green fluorescence detection. A total of 40,000 events were collected per sample, and data were analyzed using FlowJo software (BD Biosciences). Bacterial events were first identified based on forward-and side-scatter characteristics (FSC-A versus SSC-A) to exclude debris and background events. Singlet events were then selected from the gated bacterial population to exclude aggregates and doublets. Fluorescence intensity was subsequently quantified within the singlet bacterial population. The same gating strategy was applied to all samples, and a representative flow cytometry gating strategy is shown in Supplementary Figure 10.

### Intrabacterial pH measurements

For pH_IB_ measurements, an episomal plasmid harboring a streptomycin resistant cassette and allowing the expression of the ratiometric pHGFP reporter downstream the promoter Ptb38, as similarly previously described^9^, was transformed into WT, Δ*ldtB,* and Δ*ldtB*::*ldtB* strains and selected on 7H10 plates containing streptomycin. Cultures were grown to mid-log phase in Middlebrook 7H9 medium, harvested by centrifugation, and washed twice with PBS-tyloxapol 0.05%. Bacteria were diluted into Sauton’s medium adjusted to pH 5 or pH 7 in T75 flasks at a starting OD_580nm_ of 0.05. Cultures were supplemented with 200 µM oleic acid every 2-3 days to maintain growth. At designated time points, aliquots were collected for CFU enumeration and fluorescence measurements. Samples were adjusted to OD_580nm_ = 1, and 200 uL of samples were transferred into a black clear-bottom 96-well plate. Fluorescence intensities were recorded at excitation wavelengths of 395 nm and 475 nm using a Molecular Devices M5 Spectramax plate reader. The 395/475 nm fluorescence ratio was calculated and converted to intrabacterial pH values using a standard calibration curve. Three wells were used for each condition tested.

### Calcein-AM uptake assays

*M. tuberculosis* H37Rv WT, Δ*ldtB,* and complemented Δ*ldtB*::*ldtB* strains were cultured in Middlebrook 7H9 medium to mid-log phase. Cells were harvested by centrifugation, washed twice with PBS-tyloxapol 0.05%, and inoculated into 7H9 HBO medium adjusted to pH 5 or pH 7. On day 10, cultures were harvested by centrifugation, washed once with PBS-tyloxapol 0.05%, and adjusted to OD_580nm_ = 0.8 in PBS-tyloxapol 0.05% supplemented with 0.4% glucose. A 100-µL aliquot of the bacterial suspension was added to wells of a black clear-bottom 96-well plate, followed by 100 µL of 1 µg/mL calcein-acetoxymethyl ester (calcein-AM; Cayman Chemical), prediluted from a 1 mg/mL stock in PBS containing 0.05% tyloxapol and 0.4% glucose, resulting in a final calcein-AM concentration of 0.5 µg/mL. Calcein fluorescence was measured at an excitation wavelength of 495 nm and an emission wavelength of 520 nm every 2 minutes for 60 minutes using a Molecular Devices M5 Spectramax plate reader. Fluorescence values were normalized to the optical density of the final bacterial suspension and plotted at 30 min. Three wells were used for each condition tested.

### PG sample preparation and PG analysis by UPLC-MS

*M. tuberculosis* H37Rv WT, Δ*ldtB,* and complemented Δ*ldtB*::*ldtB* strains were cultured in Middlebrook 7H9 medium to mid-log phase. Cells were harvested by centrifugation, washed twice in PBS-tyloxapol 0.05%, and inoculated into 7H9 HBO medium adjusted to pH 5 or pH 7 in T175 flasks. Cultures were incubated standing at 37 °C and collected on day 8. Pellets were resuspended in 4.5 mL of 5% (w/v) SDS, heated at 80 °C for 1 h, and removed from BSL-3 containment for downstream processing. For PG isolation, samples were centrifuged (60,000 rpm, 15 min) and washed three times with distilled water. Pellets were sequentially washed with 80% acetone and 100% acetone, dried in the SpeedVac, resuspended in water, and treated with proteinase K (3 µL in 1.1 mL 100 mM Tris-HCl, pH 8) at 37 °C for 1 h. SDS was added to 1% final, samples were boiled (100 °C, 5 min), and washed once in PBS and three times in deionized water. Cell-wall material was saponified in 3 mL of 0.5% KOH in methanol with stirring at 37 °C for 4 days to hydrolyse mycolic acids. Pellets were washed twice with methanol and twice with diethyl ether. The resulting arabinogalactan-PG complex (AG–PG) was hydrolyzed with 0.2 N H_2_SO_4_ at 85 °C for 30 min and neutralised with NaCO_3_. AG was removed by washing 3-4 times with deionized water, and samples were stored at 4 °C in 100 µL H_2_O until digestion. For PG digestion, samples were incubated overnight at 37 °C with shaking in the presence of mutanolysin (20 U; Sigma, M9901-1KU). Reactions were inactivated by boiling for 5 min at 100 °C, centrifuged, and supernatants containing digested muropeptides were collected. For reduction, 15 µL 0.5 M borate buffer (pH 9) and 20 µL freshly prepared 2 M NaBH_4_ were added per 50 µL of muropeptide solution. Reactions were incubated for 20 min at room temperature, and pH was adjusted to 3-4 using 25% orthophosphoric acid.

Liquid chromatography was performed using an Acquity H-Class UPLC system (Waters) equipped with an Acquity UPLC BEH C18 column (2.1 mm × 150 mm, 130 Å pore size and 1.7 μm particle size, Waters). Separation was carried out using a linear gradient, going from 0.1% formic acid in H_2_O to 0.1% formic acid in acetonitrile. Peak identification was performed using LC-MS, and identities were assigned using retention time. Chromatograms shown are representative of three biological replicates. Relative muropeptide amount was calculated by peak integration and dividing each peak by the total area of the chromatogram. Two-way ANOVA was used to statistically compare muropeptide abundance.

### Peptidoglycan (PG) muropeptide nomenclature

PG fragments from *M. tuberculosis* were annotated using a letter–number nomenclature adapted from established conventions (Supplementary Figure 1). The letter denotes the oligomeric state (M, monomer; D, dimer; T, trimer), and the number indicates the total number of amino acids present in the peptide stem(s) attached to muramic acid residues. For cross-linked species, the first number refers to the donor stem and the second number to the acceptor stem (e.g., D44 or D33).

To indicate muramic acid modifications, the suffix “Mt” was used for monomers or dimers in which all muramic acid residues were *N*-glycolylated. In dimers containing a mixture of *N*-glycolylated and *N*-acetylated muramic acid monomers, an asterisk (*) was added to the annotation. In the absence of an “Mt” suffix or an asterisk, monomers were interpreted as containing *N*-acetylated muramic acid, and dimers were interpreted as consisting exclusively of *N*-acetylated muramic acid monomers. L,D-transpeptidase–derived 3–3 cross-links (*meso*-DAP–*meso*-DAP) and D,D-transpeptidase–derived 4–3 cross-links (D-Ala–*meso*-DAP) were distinguished based on their characteristic stem lengths and linkage patterns. The presence of anhydro-muramic acid, indicative of PG recycling, was denoted by “N” ^77^.

### MIC determinations

*M. tuberculosis* strains were grown to mid-log phase in standard Middlebrook 7H9 medium, harvested by centrifugation, and washed twice with PBS-tyloxapol 0.05%. Bacterial were diluted to an OD_580nm_ of 0.05 in 7H9 HBO medium adjusted to either pH 5 or pH 7. Aliquots of the diluted cultures were dispensed into 384-well plates in triplicate for each antibiotic concentration, along with triplicate control wells containing no antibiotic. Antibiotics were added using an HP D300e digital dispenser. Plates were incubated statically at 37°C, and bacterial growth was assessed by measuring OD_580nm_ after 14 days using a microplate reader. Meropenem, amoxicillin-clavulanate, isoniazid, and vancomycin were obtained from Sigma-Aldrich.

### Preparation of BMDMs and macrophage infection assays

Bone marrow cells were isolated from three female C57BL/6 mice aged 6-8 weeks and differentiated into BMDMs by culturing in complete DMEM supplemented with 10% heat-inactivated fetal bovine serum (FBS; Sigma), 1% HEPES (Gibco), and 30 ng/mL recombinant murine macrophage colony-stimulating factor (mCSF; Peprotech). On day 3, cultures were replenished with an additional 30 ng/mL mCSF. On day 7, macrophages were seeded into 96-well plates at a density of 8 × 10⁴ cells per well. Where indicated, macrophages were activated overnight with 20 ng/mL IFNγ (Peprotech). All animal procedures were approved by the Institutional Animal Care and Use Committee (IACUC) of Weill Cornell Medicine under protocol 0601-14A1.

For infections, *M. tuberculosis* strains were grown to early-log phase, harvested by centrifugation, washed, and resuspended in PBS-tyloxapol 0.05%. A single-cell suspension was obtained by low-speed centrifugation (800 rpm, 12 min) before dilution into DMEM. Macrophages were infected at a multiplicity of infection (MOI) of 0.1. After 4 h, cells were washed twice with warm PBS to remove extracellular bacteria. When required, meropenem was added to a final concentration of 10, 100, or 1000 μM. Intracellular bacteria were enumerated on days 0, 2, and 5 by lysing macrophages with 0.01% Triton X-100 and plating serial dilutions of lysates on Middlebrook 7H10 agar supplemented with 0.4% activated charcoal. Fresh medium containing mCSF (30 ng/mL) and the corresponding antibiotic concentrations was added to the wells on day 3 to maintain macrophage viability and drug exposure.

### Assessment of *M. tuberculosis* Localization Within Acidic Intracellular Compartments

Macrophages were prepared similarly to the what is described for the infection assays and left untreated (resting) or activated with IFN-γ prior to infection with GFP-expressing *Mycobacterium tuberculosis* (Mtb-GFP). To assess bacterial exposure to acidic intracellular environments, Mtb-GFP was labeled with pHrodo™ Red succinimidyl ester (pHrodo™ Red SE; Molecular Probes, P36600; Ex/Em 560/585 nm) as previously described^8^. The dye was resuspended according to the manufacturer’s instructions by dissolving 1 mg of pHrodo Red SE in 150 μL DMSO. Prior to labeling, a 10mL mid-log phase bacterial culture was washed and resuspended in 200 μL of 1× PBS. Bacteria were then incubated with 1 μL of pHrodo Red SE for 30 min at room temperature, washed to remove excess dye, and used for macrophage infection. Macrophages were infected at a multiplicity of infection (MOI) of 5 and incubated for 24 h. Acidification of *M. tuberculosis*-containing compartments was evaluated by fluorescence microscopy, with increased pHrodo fluorescence indicating exposure of bacteria to acidic conditions. As a supplemental and independent measure of compartment acidification, resting and IFN-γ-activated macrophages were infected with Mtb-GFP at an MOI of 5. At 23 h post-infection, cells were incubated with LysoTracker Red DND-99 (Thermo Fisher Scientific) at 1 µM final concentration for 1 h according to the manufacturer’s recommendations to label acidic intracellular compartments. Fluorescence microscopy was subsequently performed at 24 h post-infection, and the localization of Mtb-GFP relative to LysoTracker-positive compartments was assessed. For both methods, a BZ-X Series All-in-One Fluorescence Microscope (Keyence) was used on live samples (no fixation of the samples) in 96 well plates. No Z-stacks were performed to capture sequential sections along the axial (z) axis. For image analysis, the percentage of pHrodo-positive *M. tuberculosis* bacilli and the percentage of *M. tuberculosis* bacilli colocalizing with LysoTracker-positive compartments were quantified using the ImageJ software. A total of 9-10 images were analyzed per condition. Data are presented as mean ± standard deviation (SD). Statistical significance was determined using an unpaired two-tailed Student’s t-test.

## Acknowledgments

We thank Juan Jimenez and Lee Cohen-Gould for their assistance with transmission electron microscopy at the Weill Cornell Medicine Microscopy and Image Analysis Core Facility. We are also grateful to Tomas Baumgartner of the Weill Cornell Medicine Flow Cytometry Core Facility for technical support, and to Priyam Banerjee at the Rockefeller University Bio-Imaging Resource Center for imaging assistance. We thank Laura Alvarez for discussions related to PG UPLC-MS analyses and formulation. We thank Miljan Stupar for technical assistance with flow cytometry, and Natalie Thornton and Kaushik Raman for support with WGS data analysis. Schematics were created with BioRender (https://biorender.com). During the preparation of this manuscript, the authors used Microsoft 365 Copilot (for business/enterprise) in order to improve clarity of language. After using this service, the authors reviewed and edited the content as needed and take full responsibility for the content of the published article.

## Author contributions

Conceptualization: R.M., S.E., and A.G

Methodology: R.M., S.E., and A.G.

Investigation: R.M., D.R.V., K.O., A.N., C.H., T.R.I., M.P., F.C., and A.G.

Supervision: M.P., F.C., S.E., and A.G.

Writing – original draft: R.M., S.E., and A.G.

Writing – review & editing: R.M., D.R.V., K.O., A.N., C.H., T.R.I., M.P., F.C., S.E., and A.G.

## Data and materials availability

All data supporting the findings are available in the main text and Supplementary Materials. Raw sequencing data have been deposited in the NCBI Sequence Read Archive under BioProject accession PRJNA1077709. This BioProject contains both the TnSeq datasets and the whole-genome sequencing reads generated in this study, including *M. tuberculosis* H37Rv WT (BioSample accession SAMN63807355) and Δ*ldtB* (BioSample accession SAMN63807354). Strains produced in this study can be obtained from S.E. upon reasonable request and completion of a material transfer agreement, where applicable.

## Statistics and reproducibility

Generation of graphs and data analyses were performed using GraphPad Prism version 11 for macOS (GraphPad Software). Statistical analyses were performed as indicated in the figure legends.

## Supplementary Figures

**Supplementary Fig. 1:**
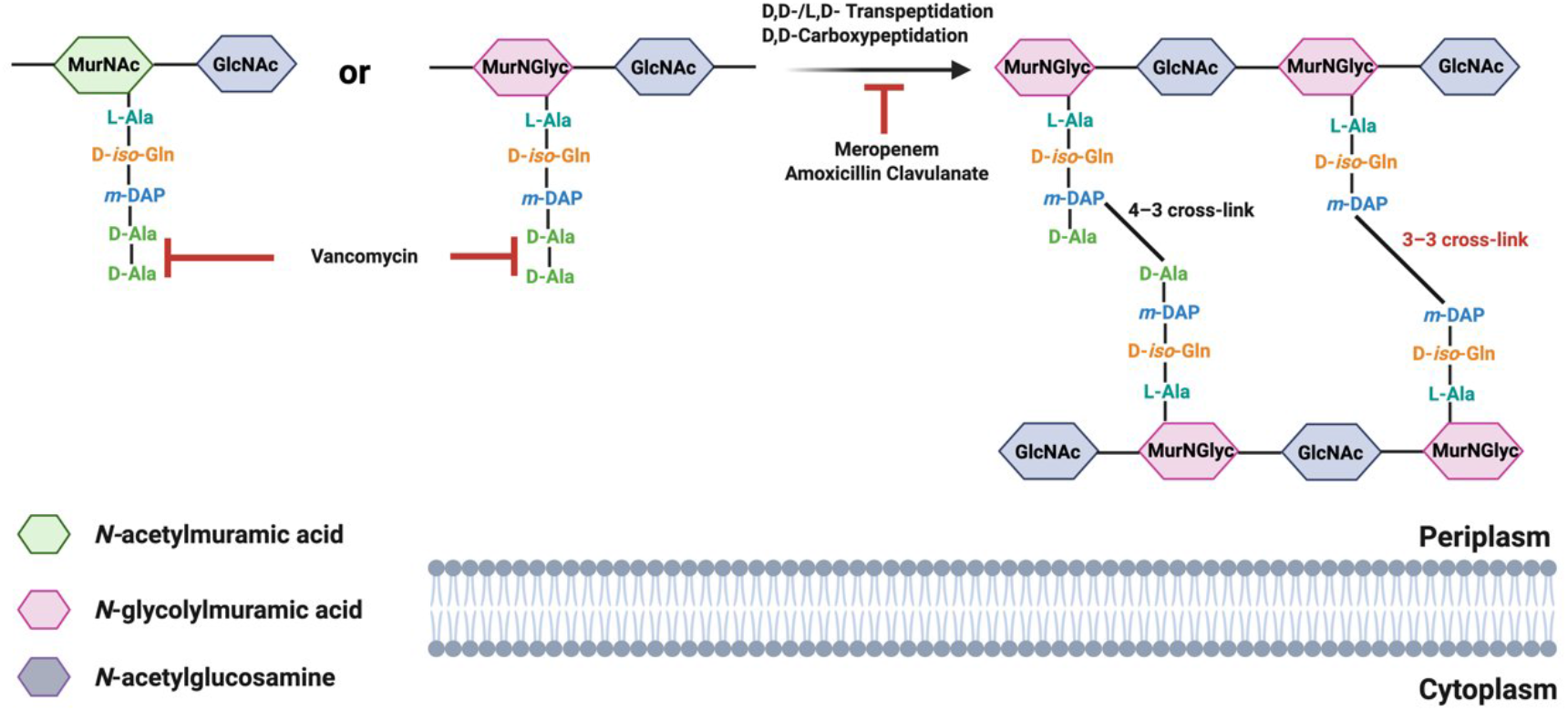
Schematic representation of *M. tuberculosis* peptidoglycan structure and the inhibition sites of PG-targeting antibiotics. Peptidoglycan (PG) subunits composed of alternating N-acetylglucosamine (GlcNAc) and either N-acetylmuramic acid (MurNAc) or N-glycolylmuramic acid (MurNGlyc) carry a conserved stem peptide (L-Ala–D-iso-Gln–meso-diaminopimelic acid [m-DAP]–D-Ala–D-Ala). In the periplasm, stem peptides are cross-linked either via D,D-transpeptidation to form 4–3 cross-links or via L,D-transpeptidation to form 3–3 cross-links. DDTs cleave the terminal D-alanine residue from the donor stem to form a cross-link between the fourth amino acid (D-Ala) of the donor stem and the third amino acid (meso-diaminopimelate, meso-DAP) of the acceptor stem. LDTs cleave the terminal (fourth) D-alanine from the donor stem and catalyze 3–3 cross-linking by joining the meso-DAP residues at the third position of both the donor and acceptor peptide stems. D,D-carboxypeptidation removes the terminal D-Ala to generate tetrapeptide substrates for L,D-transpeptidases. β-lactam antibiotics such as meropenem and amoxicillin clavulanate inhibit transpeptidation and D,D-carboxypeptidation, whereas vancomycin binds the D-Ala–D-Ala motif of pentapeptide stems, blocking downstream cross-link formation. Colored symbols denote individual sugar residues and amino acids as indicated.

**Supplementary Fig. 2:**
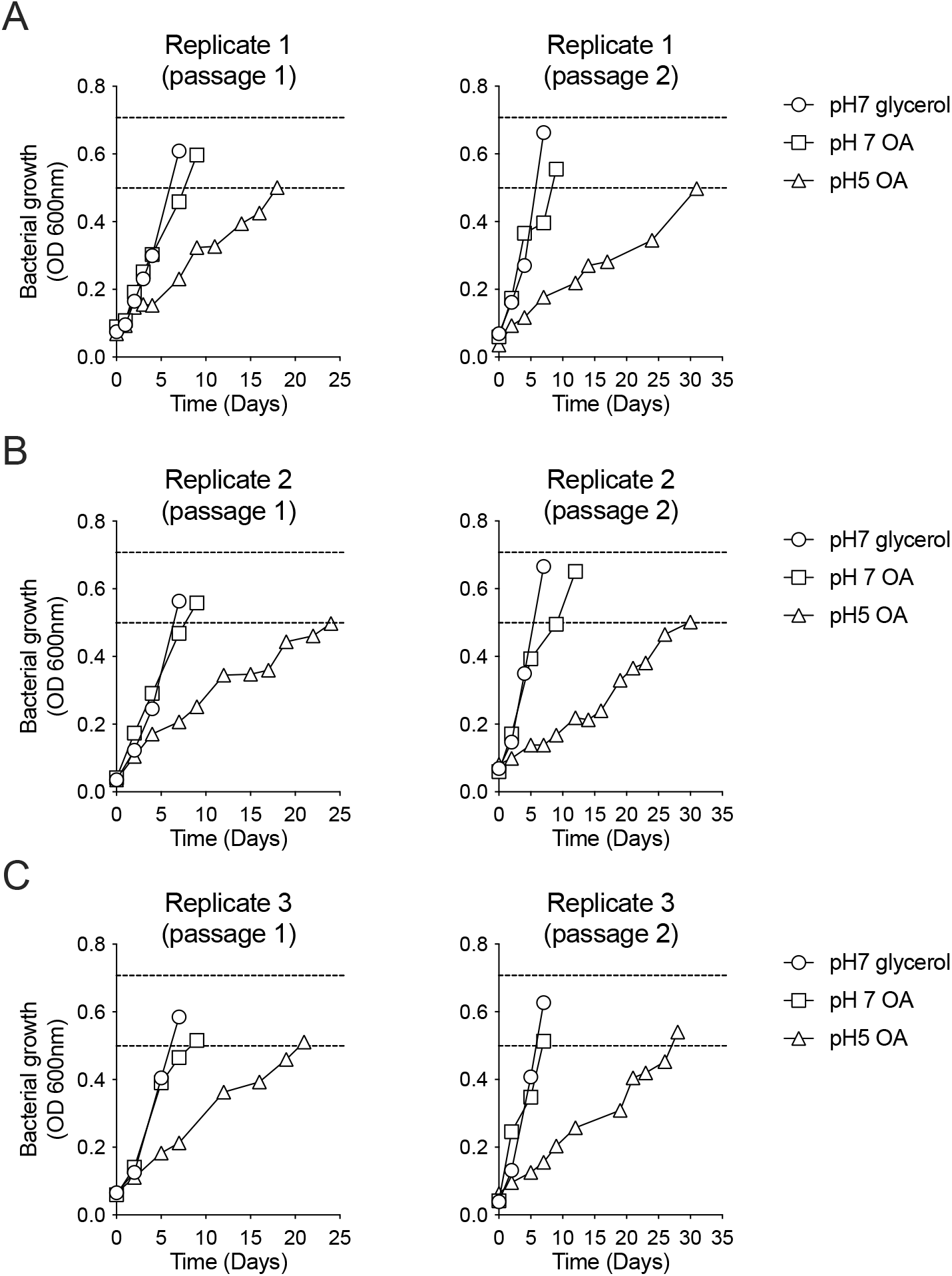
Growth patterns of the *M. tuberculosis* H37Rv transposon library under different carbon sources and pH conditions. **(A)** Growth curves for replicate 1 after first (left) and second passage (right). **(B)** Growth curves for replicate 2 after first (left) and second passage (right). **(C)** Growth curves for replicate 3 after first (left) and second passage (right). For all replicates and passages, bacteria were harvested in mid-log phase (OD= 0.5-0.7).

**Supplementary Fig. 3:**
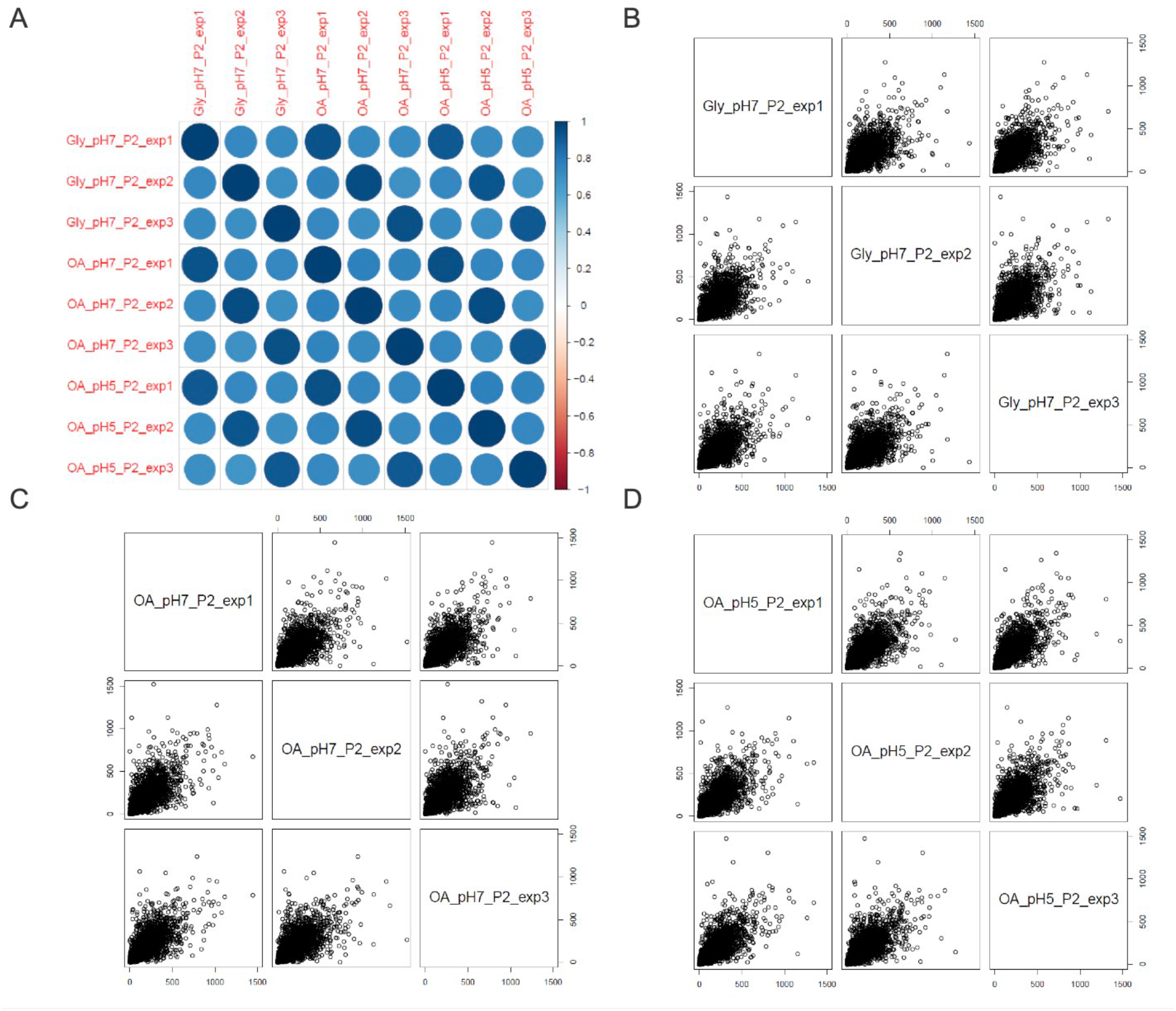
Correlation analysis of Tn-seq replicates. **(A)** Pairwise correlations (r) of gene-level mean insertion counts among samples. **(B–D)** Scatter plots of gene-level mean insertion counts across three biological replicates for glycerol at pH 7 (B), oleate at pH 7 (C), and oleate at pH 5 (D).

**Supplementary Fig. 4:**
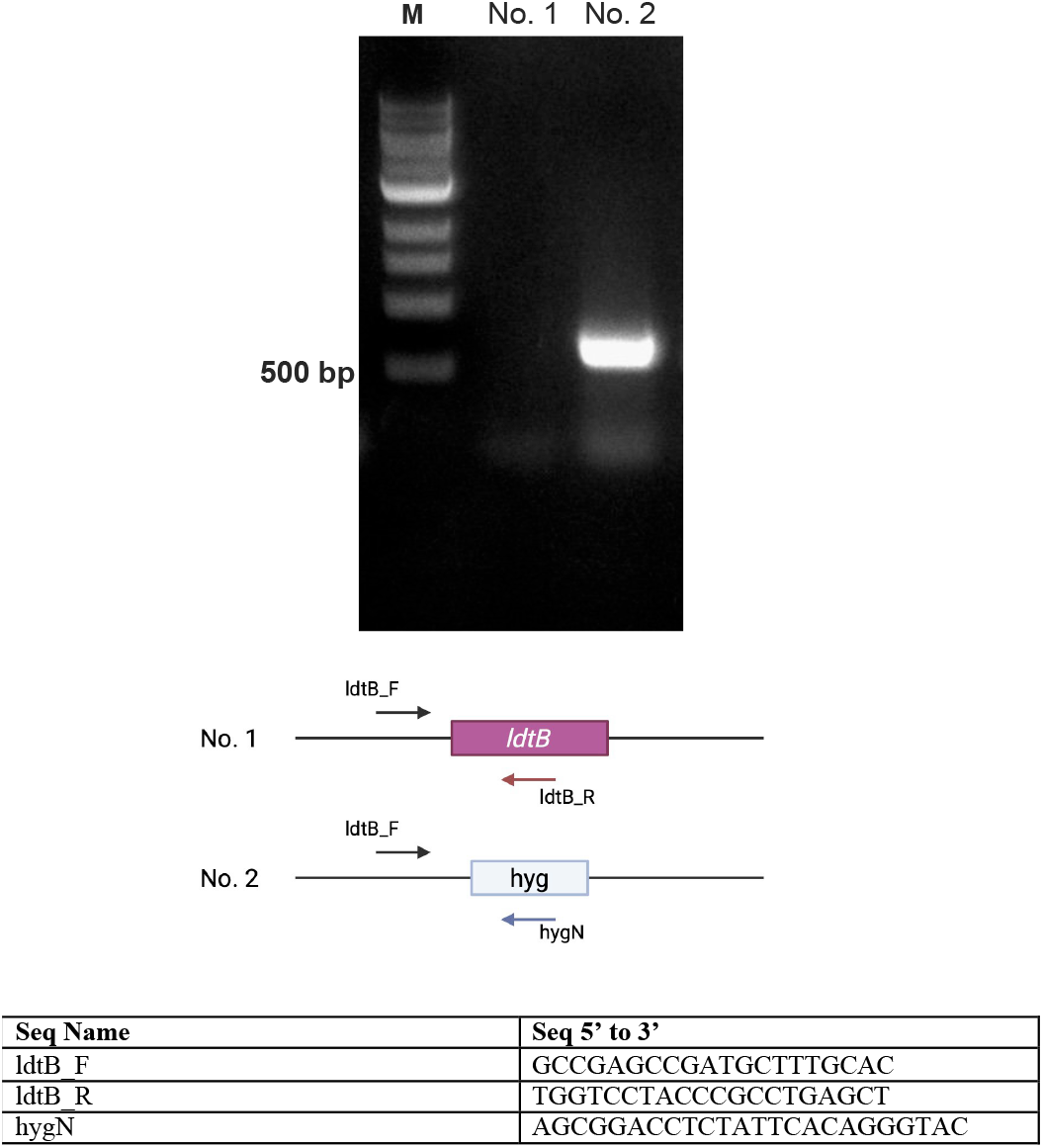
PCR verification of the Δ*ldtB* mutant. Two primer pairs were used for PCR verification. The ldtB_F primer anneals upstream of ldtB, ldtB_R anneals within the ldtB gene, and hygN anneals within the hygromycin-resistance cassette. The absence of the ldtB-specific PCR product and the presence of the hygromycin-cassette junction product confirmed deletion of *ldtB* and insertion of the hygromycin cassette.

**Supplementary Fig. 5:**
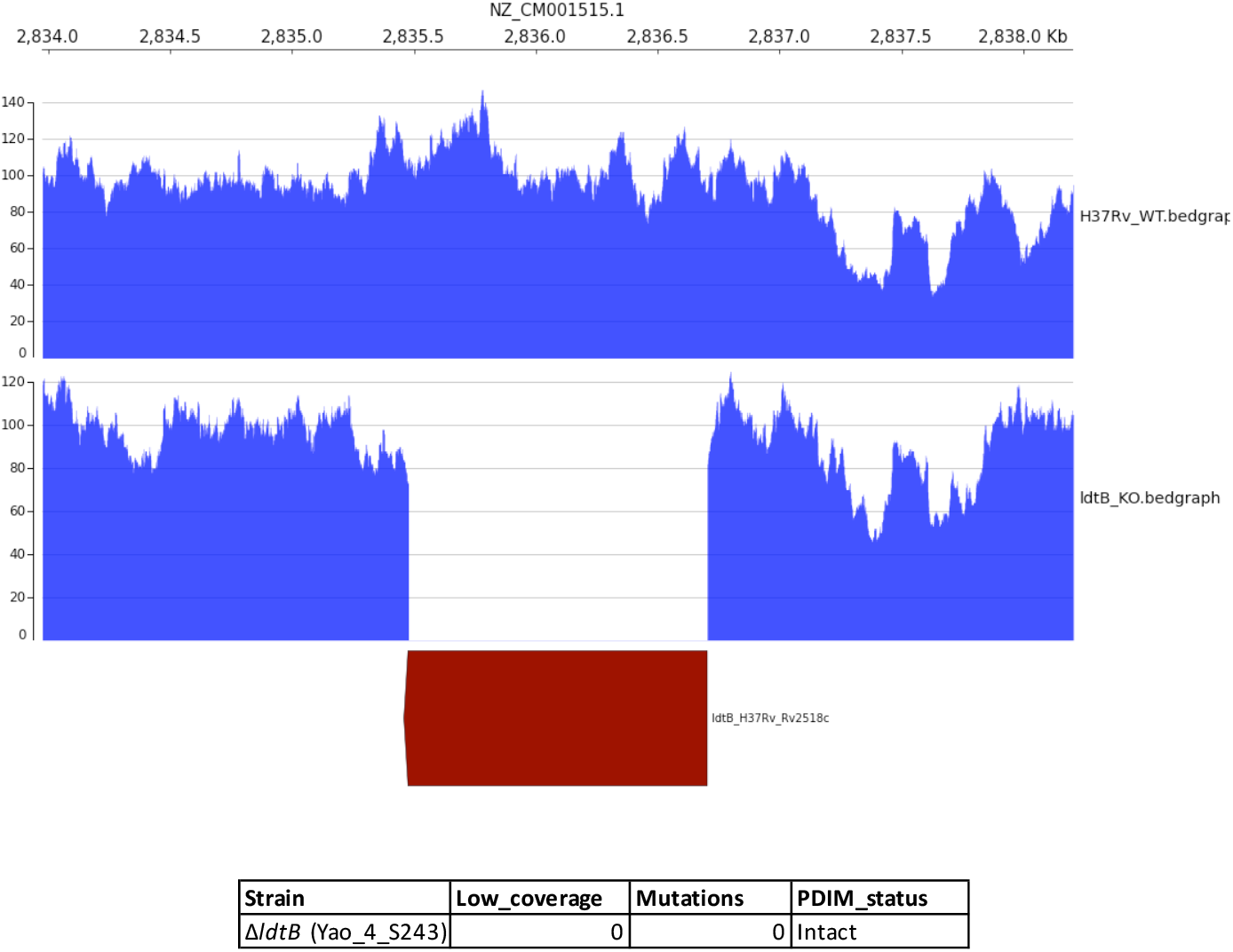
Whole-genome sequencing validation of the Δ*ldtB* mutant. Sequencing coverage of the *ldtB* locus in WT and Δ*ldtB* samples was visualized using pyGenomeTracks to confirm deletion of *ldtB*. SNP-calling analysis detected no additional secondary mutations in the Δ*ldtB* strain, including no mutations in genes associated with PDIM biosynthesis or function.

**Supplementary Fig. 6:**
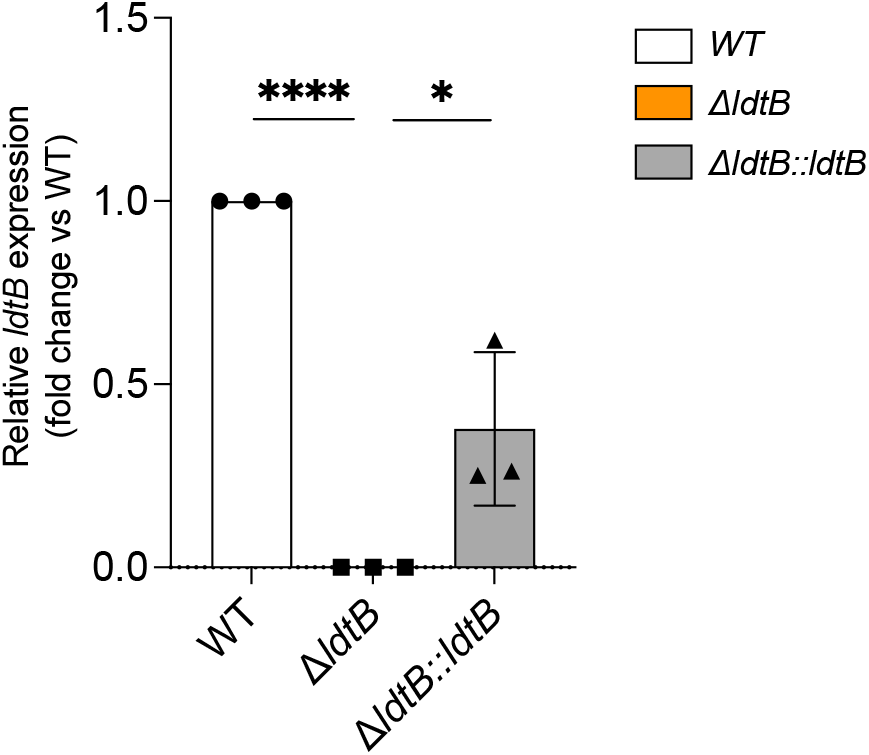
The expression of *ldtB* in WT, Δ*ldtB* and Δ*ldtB::ldtB* was assessed by RT-qPCR. Statistical significance was determined using two-way ANOVA with Dunnett’s multiple comparisons; *, adjusted *p* < 0.05; ****, adjusted *p* < 0.0001. Data represent the means of three technical replicates and are representative of two independent experiments.

**Supplementary Fig. 7:**
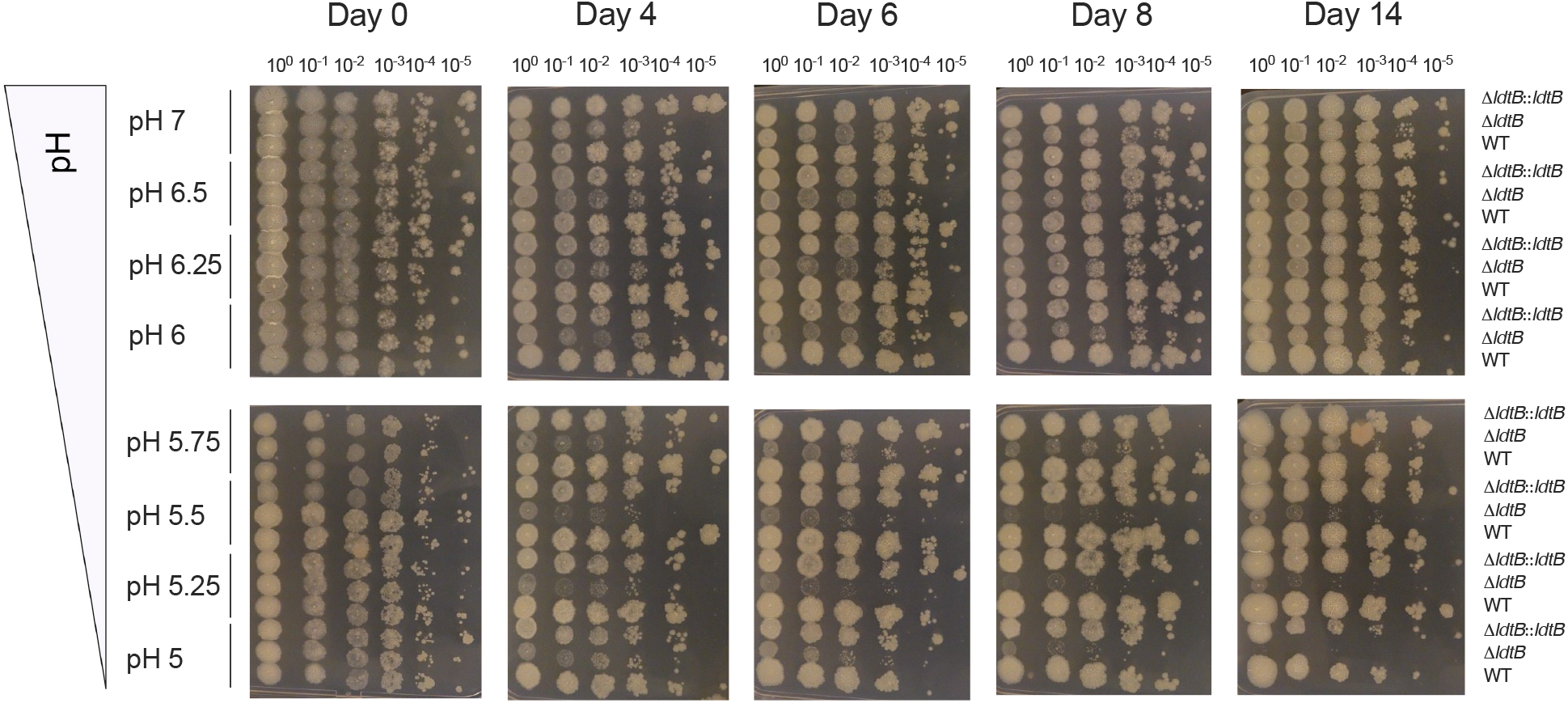
7H10 plate images of spot assays. WT, Δ*ldtB*, and Δ*ldtB::ldtB* strains were grown at various pH and serial dilutions of each culture were spotted overtime onto plates on days 0, 4, 6, 8, and 14. Data are representative of three independent experiments.

**Supplementary Fig. 8:**
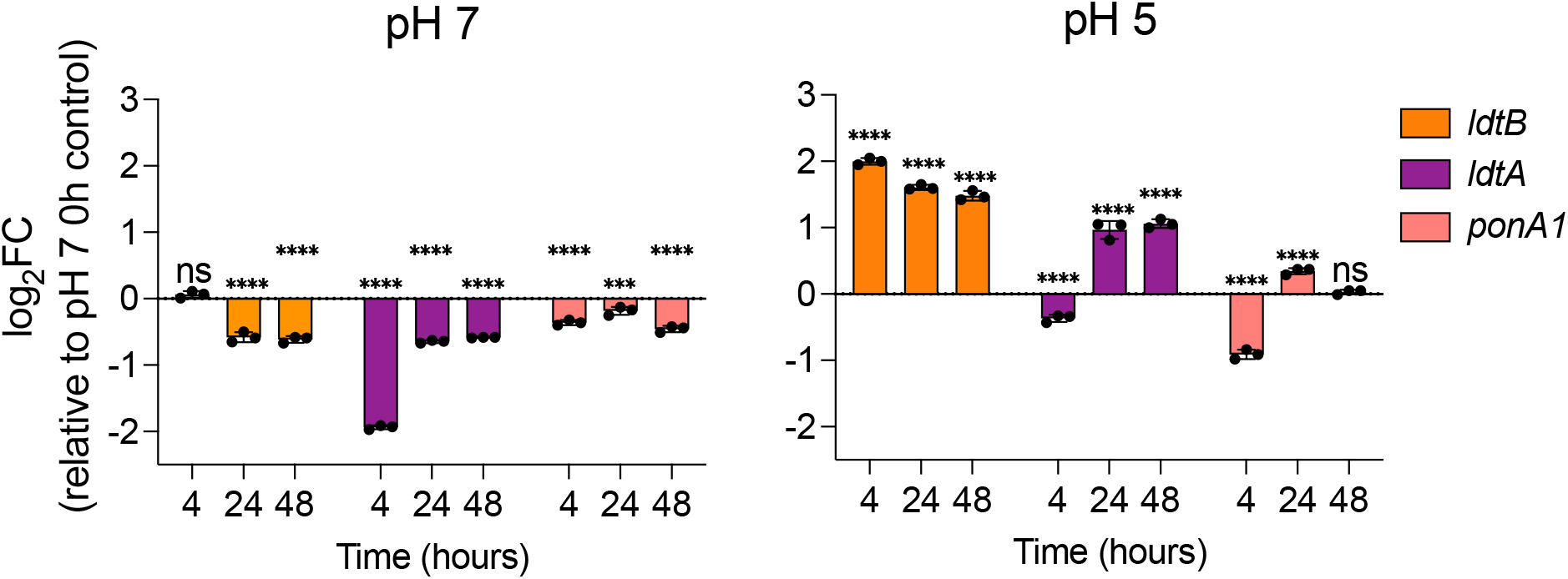
Expression of *ldtA*, *ldtB*, and *ponA1* in 7H9 medium supplemented with oleic acid (OA) at pH 7 or pH 5. Gene expression levels are presented as log₂ fold change relative to the 0 h time point at pH 7 and are shown as mean ± SD from three independent biological replicates (n = 3). Statistical significance was determined using two-way ANOVA with Dunnett’s multiple comparisons versus 0 h within each pH; ns, not significant; ***, adjusted *p* < 0.001; ****, adjusted *p* < 0.0001.

**Supplementary Fig. 9:**
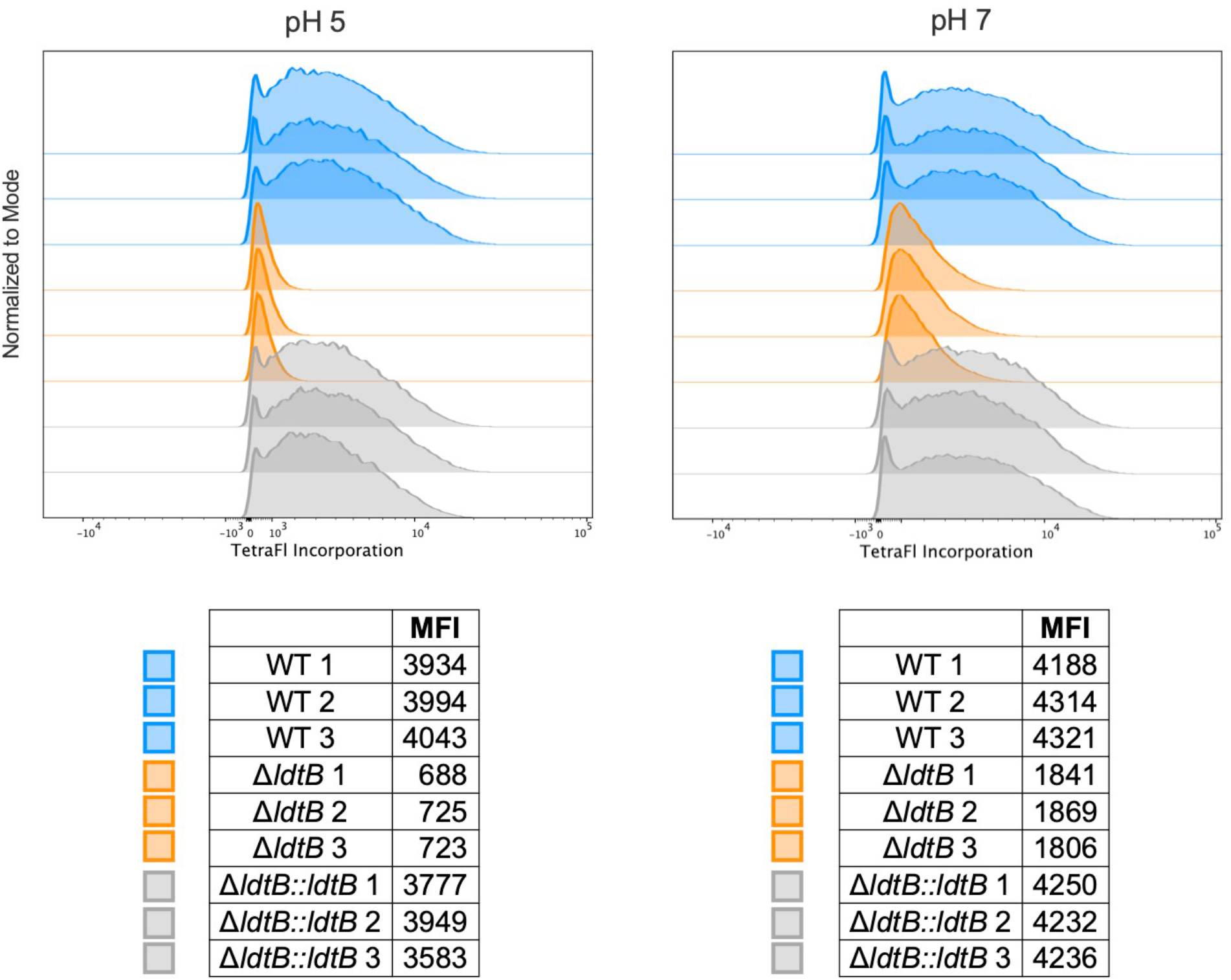
Flow cytometric analysis of TetraFI incorporation in the indicated strains at pH 5 and pH 7. Stacked histograms show the fluorescence distributions of WT, Δ*ldtB*, and Δ*ldtB::ldtB* from three technical replicates, normalized to mode. Tables show the corresponding geometric mean fluorescence intensity (MFI) values for each replicate.

**Supplementary Fig. 10:**
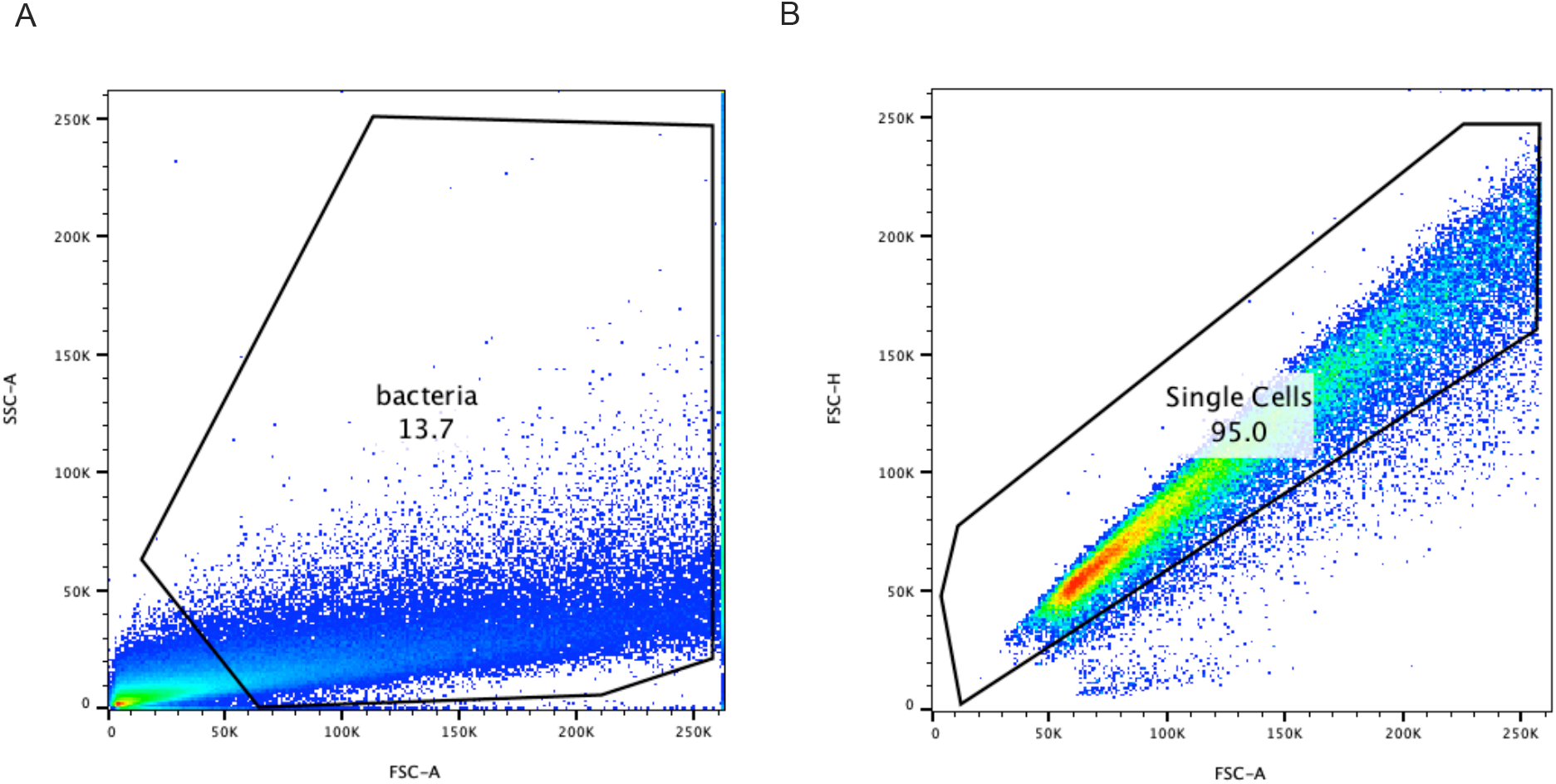
Representative flow cytometry gating strategy. **(A)** Bacterial events were identified based on FSC-A versus SSC-A to exclude debris and background events. **(B)** Singlet events were subsequently gated using FSC-A versus FSC-H to exclude aggregates. The same gating strategy was applied to all samples.

**Supplementary Fig. 11:**
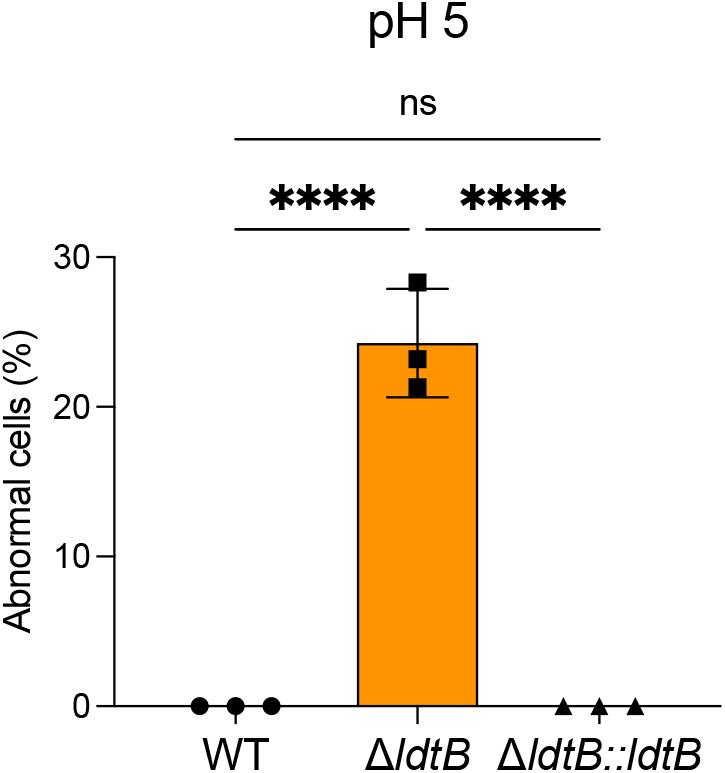
Quantification of morphologically abnormal cells in WT, Δ*ldtB*, and complemented Δ*ldtB::ldtB* strains after 8 days at pH 5 or pH 7. The percentage of abnormal cells was calculated as the number of cells displaying abnormal morphology divided by the total number of cells examined. Statistical significance was assessed by one-way ANOVA followed by Tukey’s multiple-comparisons test. ****P < 0.0001; ns, not significant.

**Supplementary Fig. 12:**
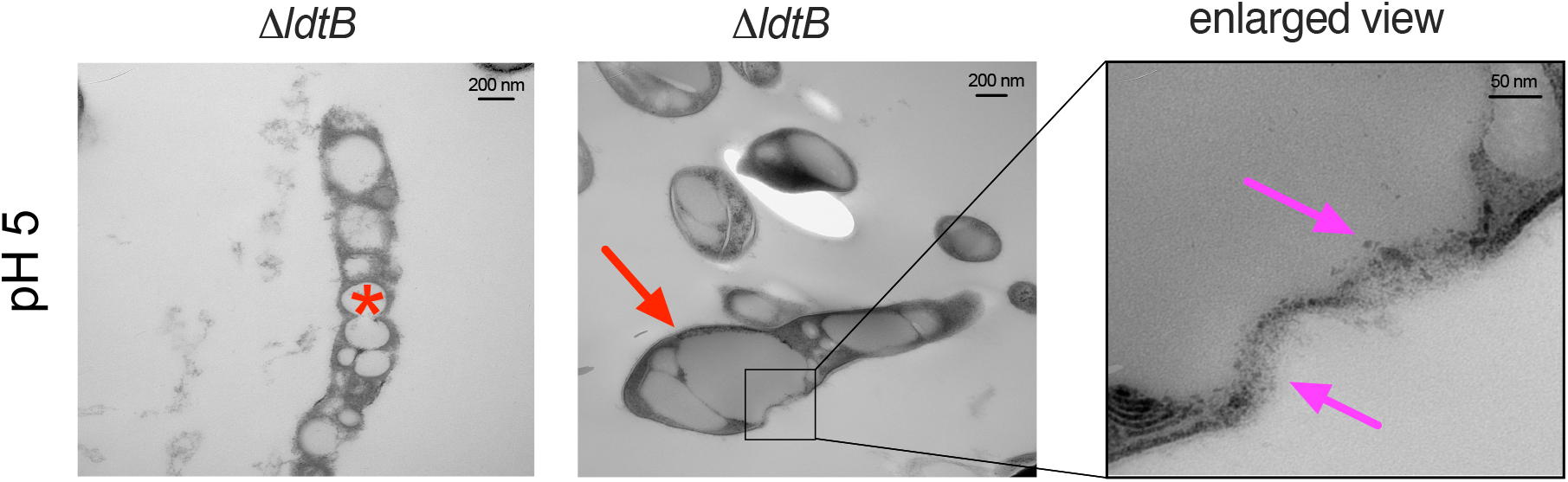
Representative transmission electron micrographs of Δ*ldtB* cells cultured at pH 5 for 8 days. Acid-specific morphological and cell envelope abnormalities in the Δ*ldtB* mutant include cellular bulging, ghost cell morphology, and localized cell envelope defects. The red arrow indicates a bulging region, the asterisk denotes a ghost cell characterized by loss of normal cytoplasmic electron density, and the purple arrows indicate localized cell envelope defects. The boxed region in the middle panel is shown at higher magnification in the right panel. Images were acquired at an accelerating voltage of 100 kV and nominal magnifications of ×50,000, ×40,000, and ×250,000 for the left, middle, and right panels, respectively. Scale bars, 200 nm in the left and middle panels and 50 nm in the right panel.

**Supplementary Fig. 13:**
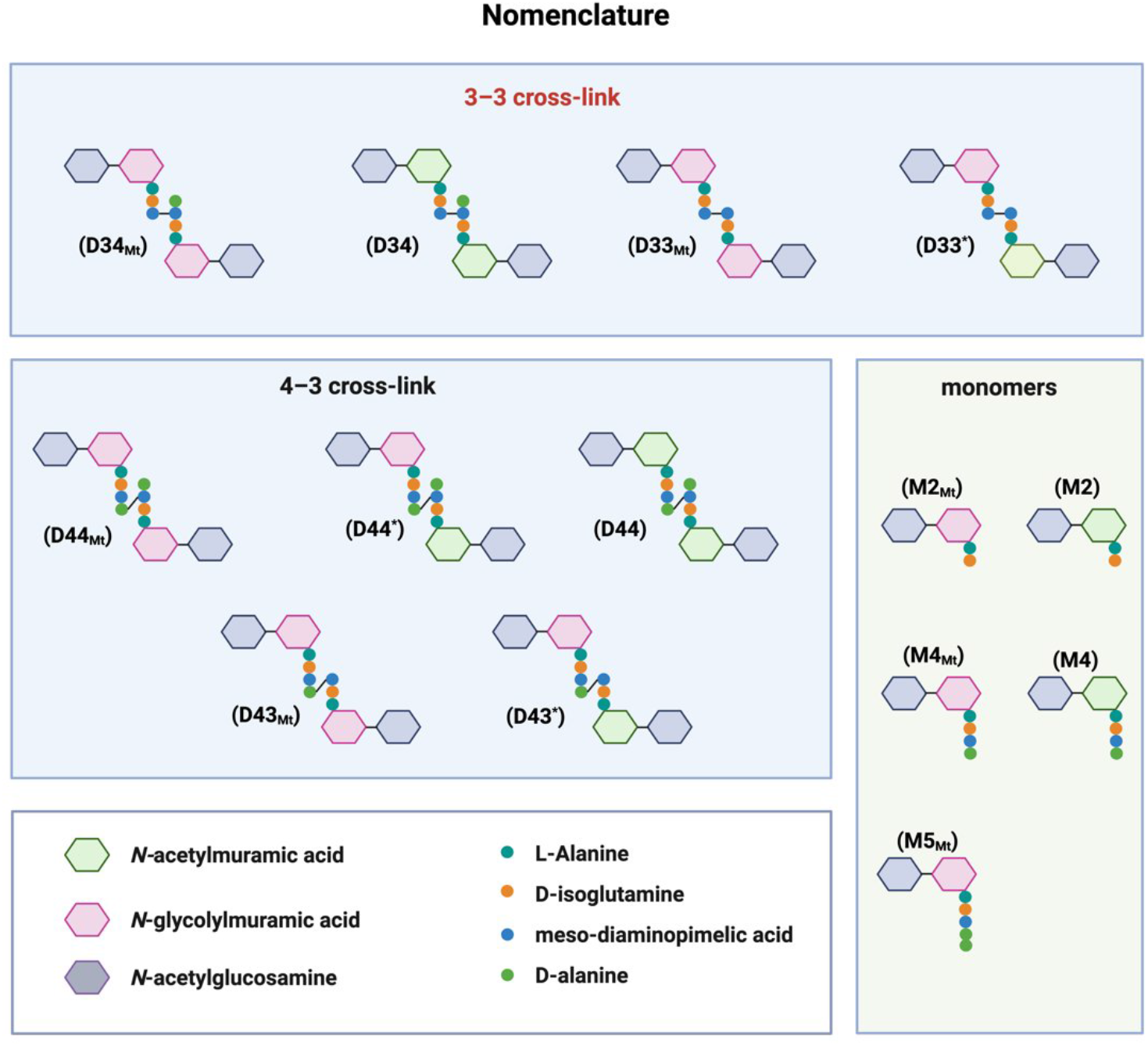
Peptidoglycan muropeptide nomenclature. PG fragments from *M. tuberculosis* were annotated using a letter–number system, where M, D, and T denote monomers, dimers, and trimers, respectively, and numbers indicate peptide stem length (donor–acceptor for cross-linked species). The suffix “Mt” denotes fully N-glycolylated species, an asterisk (*) indicates dimers containing mixed N-glycolylated and N-acetylated muramic acid, and unlabeled species are fully N-acetylated. L,D-and D,D-transpeptidase–derived 3–3 and 4–3 cross-links were distinguished by stem length and linkage chemistry, and anhydro-muramic acid was denoted by “N”.

**Supplementary Fig. 14:**
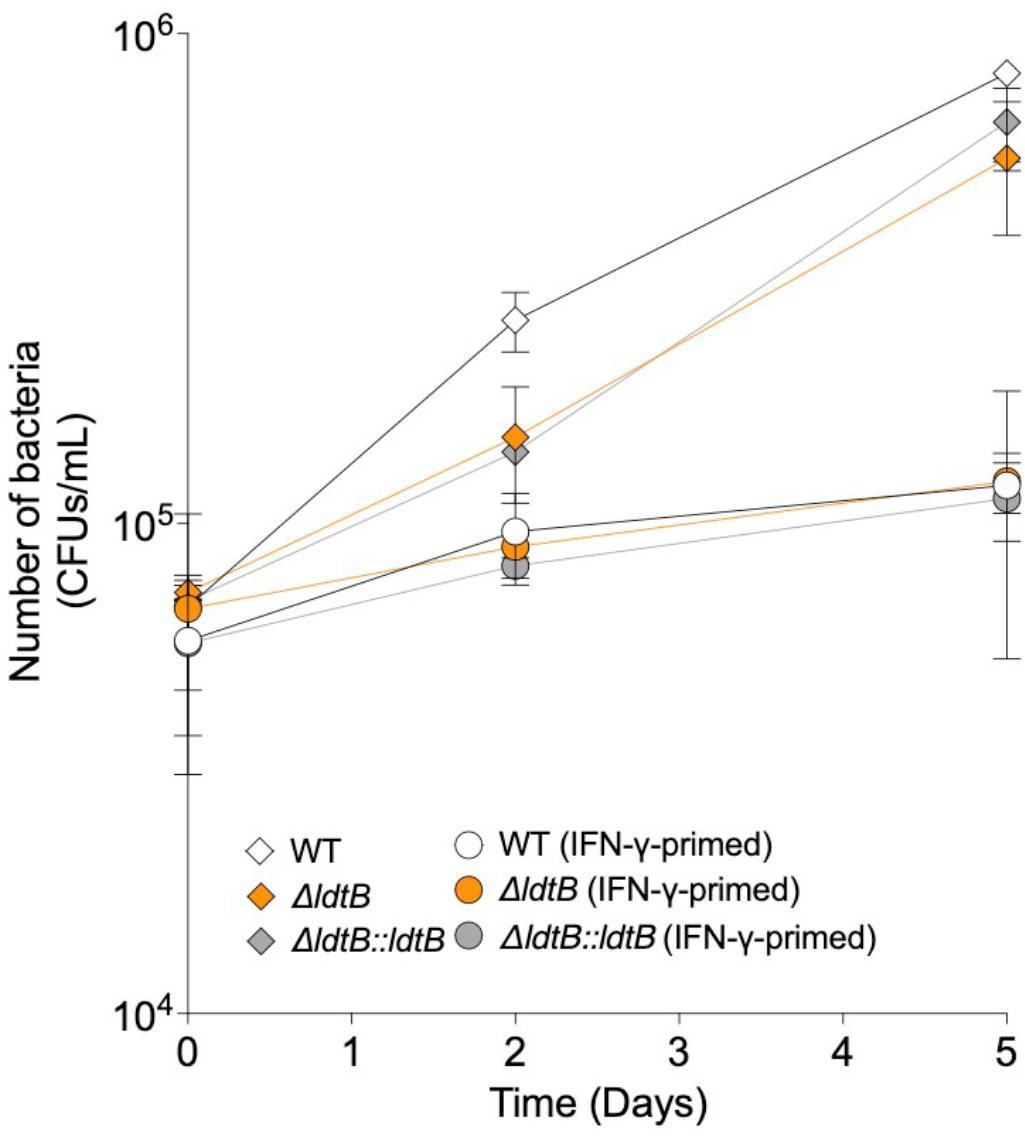
Survival of WT, Δ*ldtB*, and Δ*ldtB::ldtB* strains in BMDMs. Intracellular bacterial burden was quantified by CFU enumeration at the indicated time points over a 5-day infection period, either in the absence or presence of IFN-γ activation. Statistical significance was determined using one-way ANOVA with Tukey’s multiple comparisons test. No significant differences in intracellular survival were observed among the three strains under either condition. CFU data represent mean ± SD from three independent wells per condition and are representative of three independent experiments.

**Supplementary Fig. 15:**
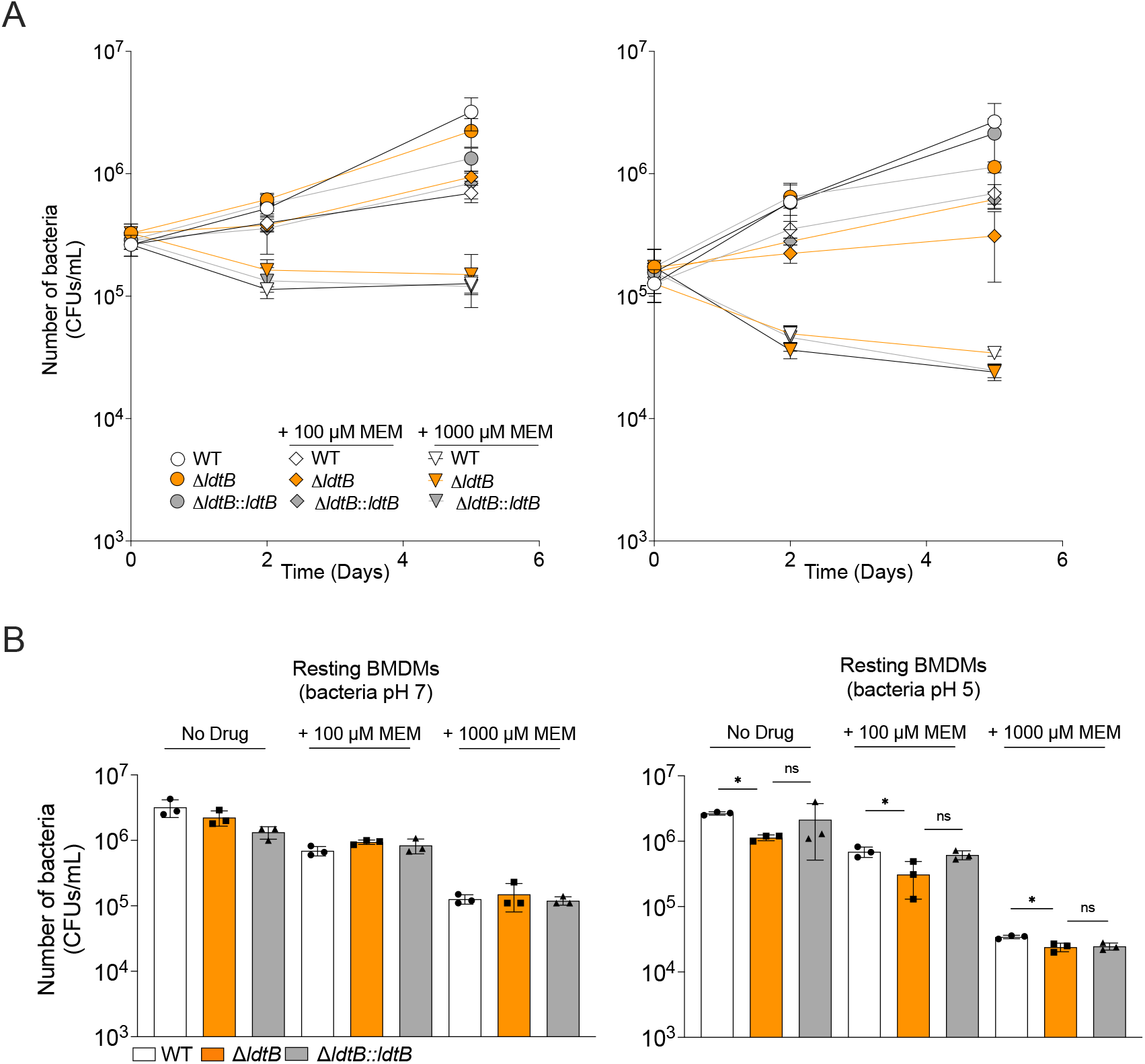
Survival of preconditioned WT, Δ*ldtB*, and Δ*ldtB::ldtB* strains in resting BMDMs treated with increasing concentrations of meropenem. Intracellular bacterial burden was quantified by CFU enumeration on days 0, 2, and 5. CFU values on day 5 were statistically compared between strains. Statistical significance was determined using one-way ANOVA with Tukey’s multiple comparisons test; ns, not significant; **, adjusted p < 0.01; ***, adjusted p < 0.001. CFU data represent mean ± SD from three independent wells per condition and are representative of three independent experiments.

**Supplementary Fig. 16:**
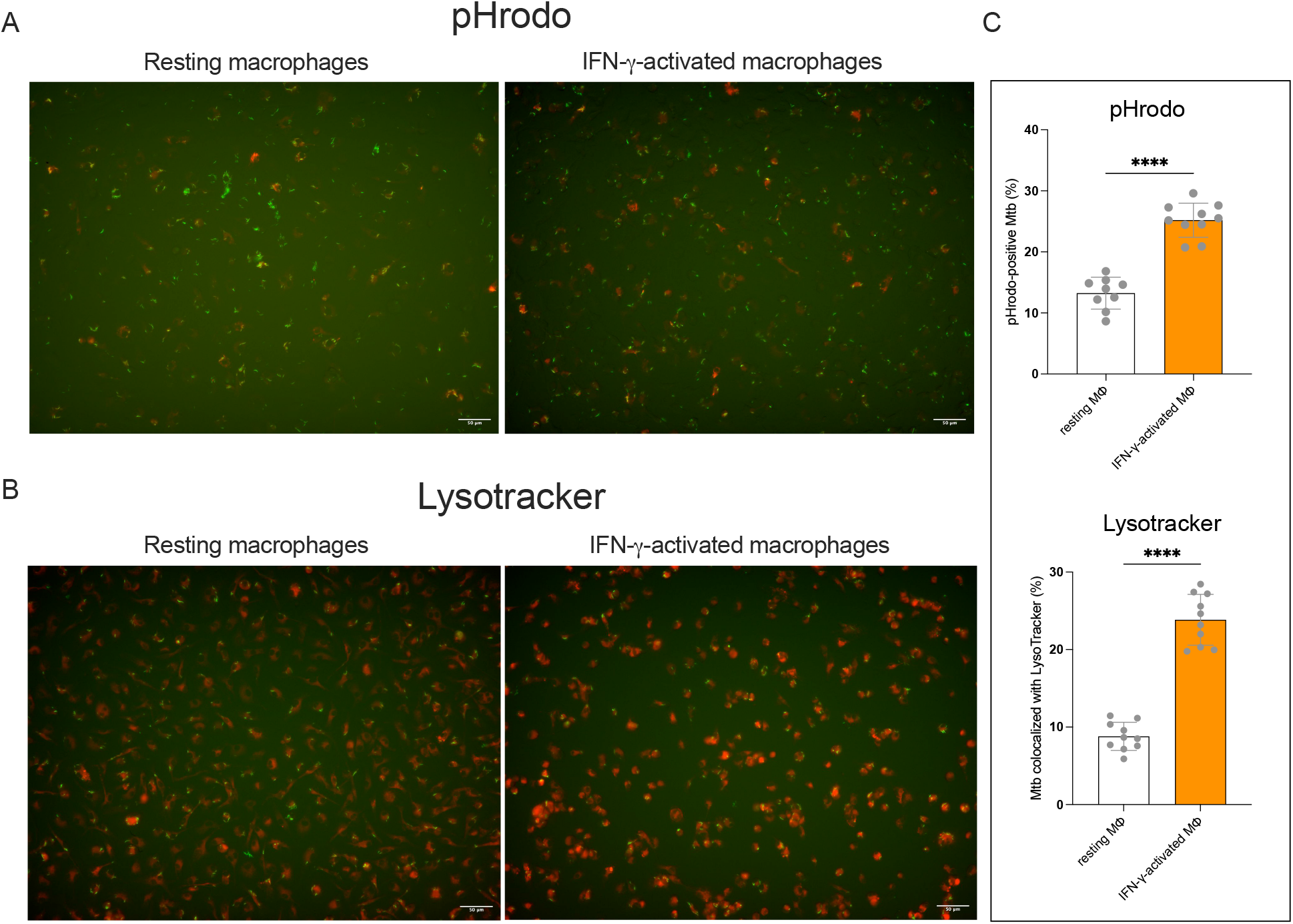
IFN-γ activation increases acidification of *M. tuberculosis*-containing intracellular compartments in macrophages. **(A)** Representative fluorescence microscopy images of resting and IFN-γ-activated macrophages infected with pHrodo-labelled *M. tuberculosis*-GFP (Mtb-GFP). Mtb-GFP was labelled with pHrodo before infection, with increased pHrodo fluorescence indicating greater exposure to acidic conditions. Scale bars, 50 μm. **(B)** Representative fluorescence microscopy images of resting and IFN-γ-activated macrophages infected with Mtb-GFP and stained with LysoTracker to label acidic intracellular compartments. LysoTracker was added 1 h before imaging. For both approaches, macrophages were infected with Mtb-GFP at an MOI of 5 and analyzed at 24 h post-infection. Scale bars, 50 μm. (**C)** Quantification of the percentage of pHrodo-positive *M. tuberculosis* (top) and the percentage of *M. tuberculosis* colocalized with LysoTracker (bottom). Each point represents an individual image; 9–10 images were analyzed per condition. Data are shown as mean ± SD. Statistical significance was determined using a t-test. ****, adjusted P < 0.0001. Both approaches showed increased association of *M. tuberculosis* with acidic intracellular compartments in IFN-γ-activated compared with resting macrophages.

---

**Supplementary Table 1:** Statistics of the TnSeq samples. ‘Mapped reads’ indicates how many reads each sample mapped to a TA site in the genome. ‘saturation’ represents the fraction of TA sites with at least 1 insertion (non-zero). ‘NZmean’ indicates the mean insertion count at non-zero TA sites. Library saturation of 0.5-0.7 (50 to 70%) is suitable for comparative TnSeq experiments.

| sample | mapped reads | saturation | NZmean |
|---|---|---|---|
| Gly_pH7_P2_exp1 | 982575 | 0.608 | 21.7 |
| Gly_pH7_P2_exp2 | 1211375 | 0.623 | 26.1 |
| Gly_pH7_P2_exp3 | 609726 | 0.576 | 14.2 |
| OA_pH7_P2_exp1 | 929447 | 0.61 | 20.4 |
| OA_pH7_P2_exp2 | 1350863 | 0.632 | 28.6 |
| OA_pH7_P2_exp3 | 579311 | 0.573 | 13.6 |
| OA_pH5_P2_exp1 | 2005746 | 0.666 | 40.4 |
| OA_pH5_P2_exp2 | 1049543 | 0.622 | 22.6 |
| OA_pH5_P2_exp3 | 627096 | 0.587 | 14.3 |

**Supplementary Table 2:**
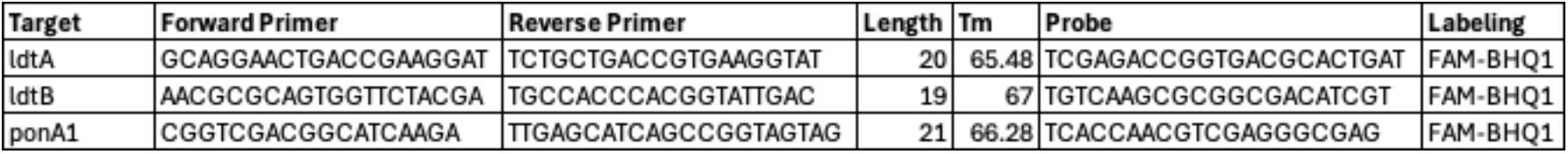
Primers and probes used for RT-qPCR.d.

